# Optimizing electrode placement and information capacity for local field potentials in cortex

**DOI:** 10.1101/2025.04.25.650658

**Authors:** Jace A. Willis, Christopher E. Wright, Ruoqian Zhu, Yilan Ruan, Joshua Stallings, Amada M. Abrego, Takfarinas Medani, Promit Moitra, John C. Mosher, Arjun Ramakrishnan, Charles E. Schroeder, Anand A. Joshi, Richard M. Leahy, Nitin Tandon, John P. Seymour

## Abstract

Recent neurosurgery advancements include improved stereotactic targeting and increased density and specificity of electrophysiological evaluation. This study introduces a subject-specific, in silico modeling tool for optimizing electrode placement and maximizing coverage with a variety of devices. The basis for optimization is the Shannon-Hartley information capacity of field potentials derived from dipolar sources. The approach integrates subject-specific MRI data with finite element modeling (FEM) used to simulate the sensitivity of subdural and intracortical devices. Sensitivity maps, or lead fields, from these models enable the comparison of different electrode placements, contact sizes, contact configurations, and substrate properties, which are often overlooked factors. One key tool is a genetic algorithm that optimizes electrode placement by maximizing information capacity. Another is a sparse sensor method, Sparse Electrode Placement for Input Optimization (SEPIO), that selects the best sensor subsets for accurate source classification. We demonstrate several use cases for clinicians, engineers, and researchers. Overall, these open-source tools provide a quantitative framework to select devices from a neurosurgical armament and to optimize device and contact placement. Using these tools may help refine electrode coverage with low channel count devices while minimizing the burden of invasive surgery. The study demonstrates that optimized electrode placement significantly improves the information capacity and signal quality of local field potential (LFP) recordings. The tools developed offer a valuable approach for refining neurosurgical techniques and enhancing the design of neural implants.

**Highlights:**

- tool for simulating subject-specific local field potentials and electrode sensitivity.
- ptimized electrode placement enhances ROI source coverage, and signal quality.
- sensor-based classification boosts data quality without extra electrode cost.
- comparisons of devices and contact arrangements.

**Graphical Abstract:** 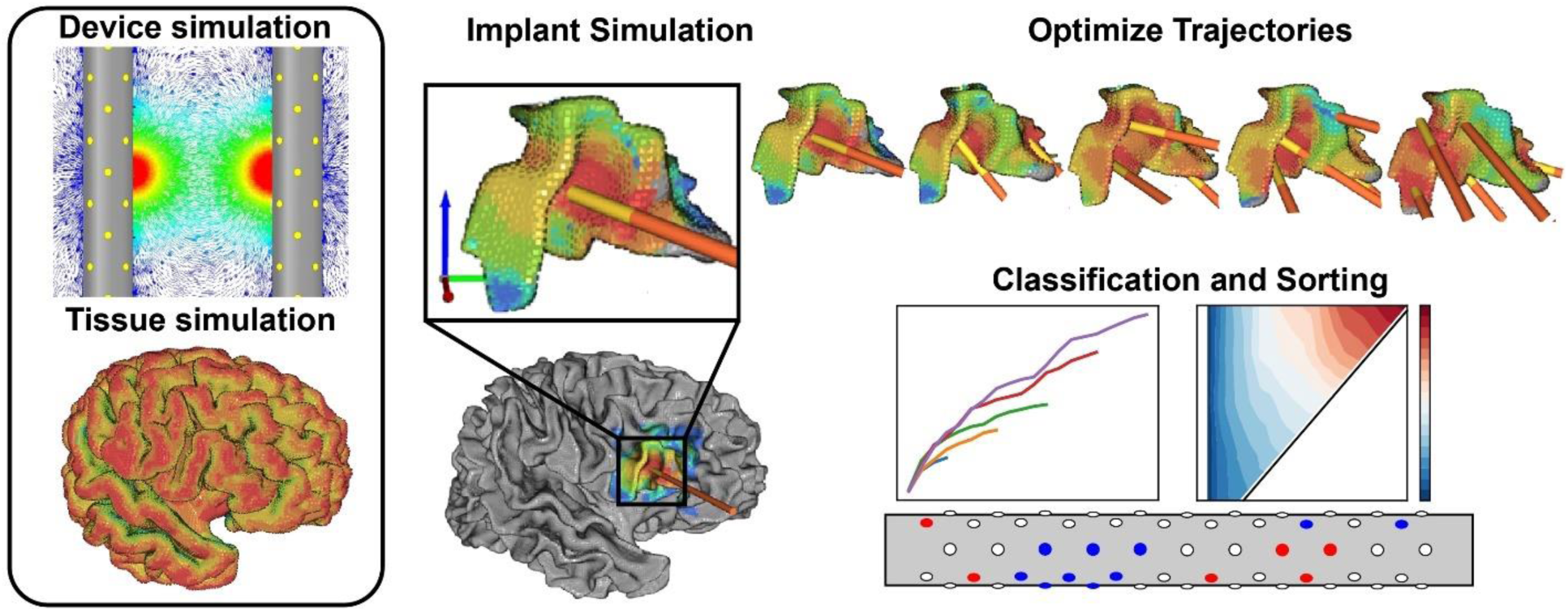

## 1. Introduction

Brain activity produces numerous biophysical information patterns, including broadband field potentials, high gamma activity, multi-unit activity, and single-unit activity that researchers have mined for decades. Each recording technology will vary in its peak signal-to-noise ratio (SNR) depending on interdependent factors that include the electrode diameter, material, substrate properties, and tissue conductivity. When expressed in units of bits, the information capacity of a given channel or sensor is related to the log base 2 of SNR, but recording arrays often contain redundant information, which is a function of the spatial extent of each biophysical source, device placement, and electrode array configuration. Device optimization can be guided by estimating the Shannon-Hartley information capacity for any given array of sensors in brain source space. A capacity measure, based on SNR and bandwidth (Equation 1), could provide a unified framework to compare and improve device performance.

Neural implants are critical tools for both neuroscience research and clinical treatment. Device placement generally follows two principles: (i) minimize anatomical and functional disruption, and (ii) maximize the SNR captured from the region of interest (ROI). These goals can conflict. In the 2000s, U.S. clinicians favored electrocorticography (ECoG) for epilepsy monitoring via craniotomy. However, subsequent studies found that stereotactic EEG (SEEG), inserted through a twist-drill hole, offered better safety and improved seizure signal quality (Bernabei et al., 2021; Tandon et al., 2019). SEEG technology has since advanced, with higher channel counts and options for single-unit recordings (Despouy et al., 2020; Fried et al., 1999). These improvements, driven by decades of refinement (Engel, 2018), have enhanced seizure localization and treatment (Bartolomei et al., 2017). At the same time, intracranial recordings are increasingly used in brain-computer interfaces (BCIs) to assist patients with paralysis or locked-in syndrome (Patrick-Krueger et al., 2024; Silva et al., 2024). To support such applications, surgical planning that incorporates subject-specific ROIs may inform better device selection and placement.

As neuroelectronic technologies evolve, quantitative tools are increasingly needed to compare existing and proposed devices in silico. Biophysical models have become more accurate and accessible through advances in computational neuroscience (Bhalla et al., n.d.; Markram et al., 2015; Medani et al., 2023; Piastra et al., 2021). For instance, Piastra et al. (2021) quantified SNR tradeoffs between EEG and MEG using models of field potentials (FPs). Similar strategies can inform intracranial device development, particularly for mesoscale recordings (∼0.5–2 mm). Much effort in neuroengineering has focused on miniaturization and high-channel-count arrays to record action potentials (Seymour et al., 2017), such as the 900-contact Neuropixels probe (Steinmetz et al., 2018). While ideal for local network recordings, these arrays are not optimized for meso- or macro-scale activity. In clinical settings, SEEG remains the standard for FP measurement, with typical designs using 8 ring contacts (2 mm length, 3 mm pitch) on a 0.8 mm diameter probe. Variants with segmented rings are emerging for deep brain stimulation (Schüpbach et al., 2017). Our lab recently developed a directional SEEG device, DiSc, which provides enhanced spatial resolution at the mesoscale (Abrego et al., 2023). Still, future devices will offer diverse tradeoffs that would benefit from a detailed pre-surgical evaluation.

Choosing an optimal device configuration for a given ROI is challenging, both for humans and algorithms, especially given limited preoperative data. Surgical planning typically emphasizes anatomical safety and hypothesis testing such as identifying seizure focal onset zones (Cardinale et al., 2013), but methods to quantitatively optimize device selection and placement are lacking. Despite the availability of biophysical models, they are rarely incorporated into planning workflows.

We propose a five-step *in silico* pipeline (Figure 1) to address this gap that focuses on FPs and derivative local FPs and should be expandable to other biophysical information sources. The process begins with computing a device-specific lead field via finite element modeling (FEM), followed by assigning dipole sources based on MRI-defined ROIs. Simulations yield voltage recordings for each sensor, enabling device capacity estimation via the Shannon-Hartley theorem (Shannon, 1948). A genetic algorithm then optimizes trajectories by maximizing total capacity over the ROI. Sparse Electrode Placement for Input Optimization (SEPIO), adapted from Brunton et al. (2016), identifies compact, high-performing sensor subsets using PCA space to reduce computational load while preserving source classification quality. SEPIO also flags redundant contacts, supporting streamlined device designs. To validate our *in silico* results, we implemented a physical phantom: a PBS-filled head model with 11 dipoles simulating a central sulcus. Signals recorded by two DiSc arrays were compared to model predictions, confirming the degree of model match and the method’s classification accuracy.

**Figure 1.**
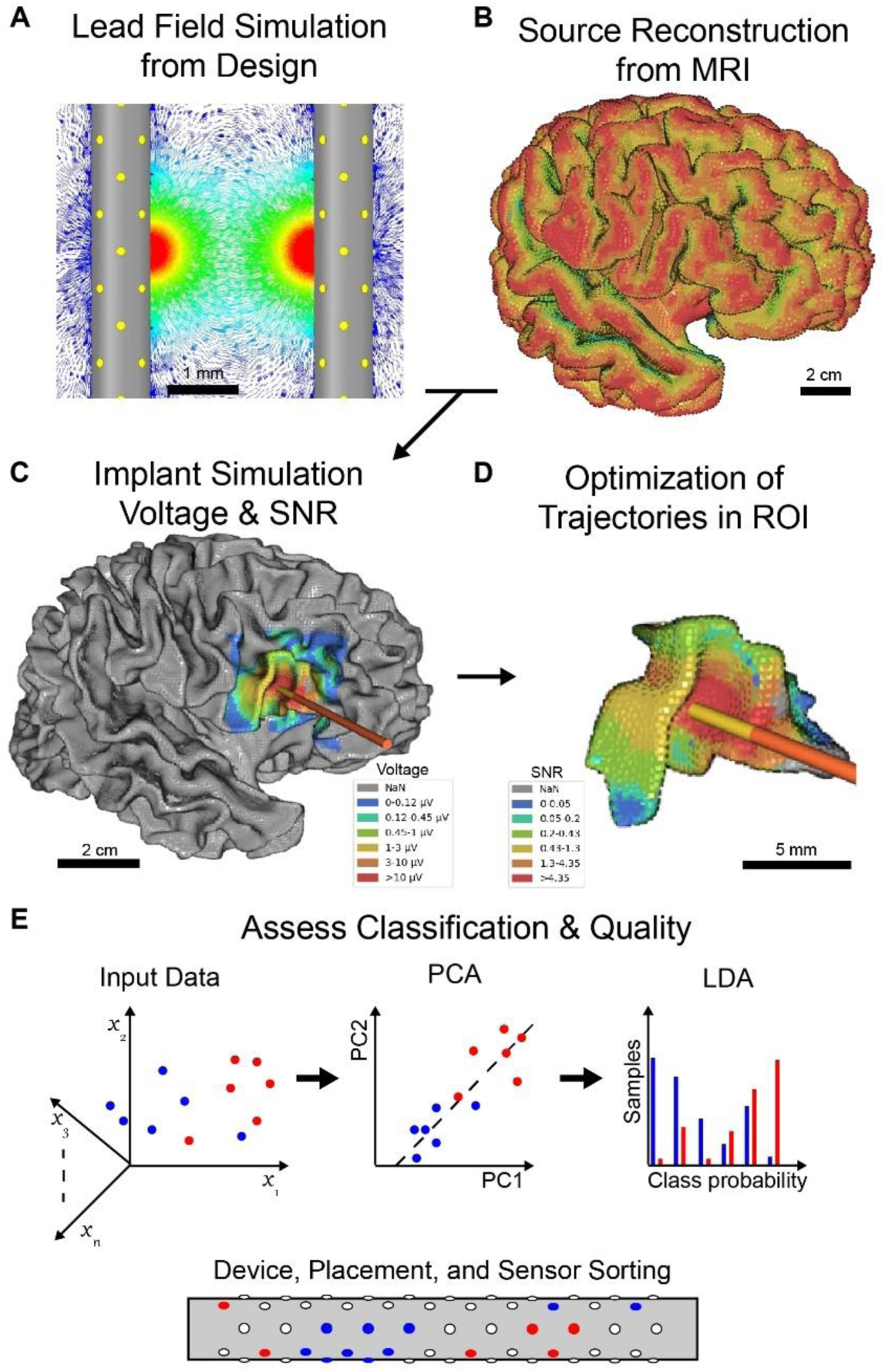
Multi-device trajectory and sensor simulation and optimization method. (A) Individual devices are digitally constructed and simulated to produce lead fields. An inner cortical surface (ICS) is extracted from T1 MRI data using BrainSuite to recreate a subject-specific source space. The ICS is inflated to form the (B) pial surface for visualization. (C) For a given device, a voltage and SNR map can be computed. (D) Isolating an ROI can greatly reduce the computation of an optimal trajectory, defined as maximum total information capacity. (E) For a given device configuration, one can use classification methods as an additional comparison of solution quality and to rank the contribution of devices and contacts, which we refer to as Sparse Electrode Placement for Input Optimization (SEPIO).

## 2. Methods

Engineers, clinicians, and neuroscientists vary on some specific vocabulary that we will define here. Electrode, device, and implant all refer to a single whole device, while sensor, contact, and channel refer to a single recording surface. Devices placed inside the cortex are referred to as both depth electrodes and probes, while subdural devices may be referred to as surface electrodes, or grids. Voltages are the primary data and calculated for every source and sensor pair. The manner in which voltages are used may change the ideal representation of this array. Data can be projected onto the sensors (sensor-space or sensor domain), or data can be projected onto the source surfaces (source-space or source domain).

### 2.1. Information capacity, lead field, and sensor-specific voltage calculation

The Shannon-Hartley information capacity describes the maximum information capacity to be a relationship of the bandwidth, signal power, and noise power for a given source and device (Shannon, 1948). This theorem is a fundamental principle in information theory, quantifying the theoretical upper limit for reliable data transmission on a known communication channel. Historically, the theorem served as a backbone of communications infrastructure design. Here, we apply the same principles to optimize the potential recording capacity for neural implants.

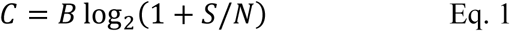

Where 𝐶 is the information capacity (bits/s), 𝐵 is the sampling bandwidth (Hz), 𝑆 is the signal power, and 𝑁 is the noise power. Signal power depends on the sensor and dipole spatial configuration. In simulation, B is defined as the Nyquist bandwidth (half of the device’s backend sampling rate). Noise N is modeled as a zero-mean normal distribution with variance specific to the device type and recording system. Using DiSc, the noise standard deviation is 4.1 𝜇𝑉, based on prior RMS noise measurements (Abrego et al., 2023).

Current dipole dynamics can be replayed in a volume conduction model having different electrode geometries and insulating properties. Lead fields capture the sensitivity of each unique sensor-source pair, termed a gain vector for a specific point in space (Baillet et al., 2001). Ansys Electronics Desktop (AEDT v.2024 R2) is used to solve lead fields by applying 1 Amp (monopolar) to each channel sequentially and recording the electric field at all elements of the finite element model (FEM), with the region boundary defined as at 0 V. Tissue around the device is defined with isotropic conductivity of 0.25 𝑆/𝑚. This method is also capable of generating lead fields based on inhomogeneous and anisotropic conductivity due to white matter, CSF, meninges, and skull from MRI or a standard head model. As discussed at length in §4.1, this is most practical with a fixed device placement. Electric field vectors are exported and averaged to a cartesian grid on which dipole currents can be defined and produce a simulated voltage on each contact *via* voltage-current reciprocity (Rush & Driscoll, 1969). The simplest form for the measured voltage (𝑉) on a sensor arising from a current dipole (𝑝⃗) at a given location (𝑟) and the lead field vector (𝐿⃗) associated with that point in space, is given by the following for each sensor index (𝑖).

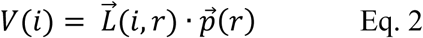

When optimizing trajectories within an ROI, source domain information capacity is used to assess the coverage of each individual source and give visual feedback of performance. After device placement is determined, sensor domain voltage is provided to SEPIO so that final optimization can be made with respect to the devices. Though classification accuracy is the primary metric, information capacity ranked by sensor priority may further evaluate the diversity of each multi-device solution.

The current dipole is our assumed model as explained in the following section. Using the recorded voltage patterns, signal coverage qualities and classification accuracy of each arrangement can be tested. A library of lead fields is provided for multiple device types and variations in the public repository (§7), including variants of ECoG, SEEG, and DiSc. We additionally refer to “virtual macros,” which are averaged rings of DiSc microelectrodes that emulate SEEG macroelectrode rings.

### 2.2. Source space simulation and visualization

Current dipoles represent the peak amplitude LFP information as previously demonstrated using a quasi-electrostatic approximation of Maxwell’s equation (Hämäläinen et al., 1993; Nunez et al., 2019). Dipoles are arranged in a single layer at a middle depth in cortex. The arrangement of the current dipoles is determined by either manual assignment or processing of MRI data using BrainSuite (refer to §2.5) (Shattuck & Leahy, 2002). The inner cortical surface (ICS) is extracted and projected into the pial surface (PS), respectively representing the grey-white interface and grey-pial interface, to accurately match vertices for visualization. Each surface contains surface triangles, corner vertices, and normal vectors to the surfaces. The PS is used for projecting realistic visualization while the ICS is used for a ‘deflated’ visualization to observe high curvature spaces. ICS vertices are used as dipole locations with an added offset in the direction of the nearest normal to account for the distance from the grey-white interface to the granular layer where the largest dipole contributors are most common (Ikeda et al., 2005). This study uses an estimated cortical thickness of 2.5 mm (Fischl et al., 2000) and an estimated half cortex depth to layer 4 (Palomero-Gallagher & Zilles, 2019), as these vary by region.

Once dipole positions are defined, the nearest normal vector to each original ICS vertex is used for the dipole direction. We apply 2 𝑛𝐴 · 𝑚/𝑚𝑚^2^based on the upper end of dipole amplitude observed by Murakami and Okada (Murakami & Okada, 2015). A column is defined by the voxel size of 500 𝜇𝑚 on a side with a cross-sectional area of 250 𝜇𝑚^2^, yielding a dipole strength of 0.5 𝑛𝐴 · 𝑚. With complete source space, device lead fields can be used to evaluate the individual or combined signal of dipole sources as described in Equation 2 and visualized in Figure 2. In this study, dipoles are evaluated one at a time, effectively observing them in isolation with matching magnitude, though this method does not exclude the ability to define unique source magnitudes. The approach efficiently determines the maximum achievable voltage, SNR, and information capacity of each source for any given arrangement of sensors or devices. Additionally, SEPIO attempts classification when sensor coefficients accurately capture signal diversity, and label-wise accuracy reflects the ability to classify a given source.

**Figure 2.**
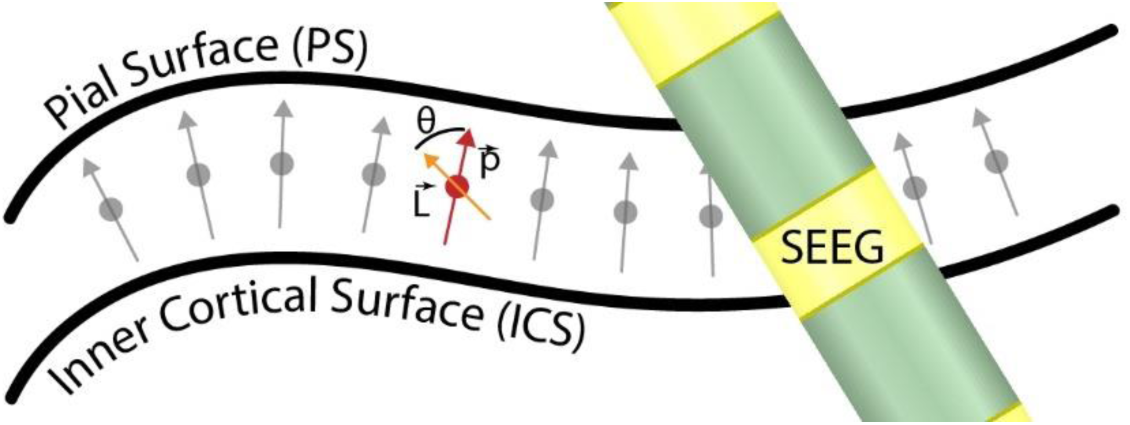
Dipole distribution is relative to the pial surface (PS) and inner cortical surface (ICS). Grey arrows represent the dipole vectors, where the midpoint is the assigned location. An SEEG device is shown in green with ring electrodes in yellow. This device and trajectory create unique gain vectors. At a given source, such as the example in red, the magnitude and angle between the source and gain vector determines the dot product scale which produces a simulated voltage (Equation 2).

Device position and trajectory are controlled by the defined source space and individual device parameters. Each device is limited to a maximum trajectory relative to scalp normal vectors and a range of permissible depth into cortex. If a device trajectory exceeded the maximum relative angle, its vector is corrected back toward the scalp normal. If a depth is outside of the allowed range, it is moved along the scalp normal vector direction to the nearest allowed depth. Additionally, a bounding cylinder is drawn around each device based on the characteristics of each. These cylinders are used to avoid unrealistic intersections and maintain a minimum buffer distance. The combination of these limiting factors allows intracortical and subdural devices to be represented adequately for simulation.

### 2.3. Multi-device trajectory optimization

Modelling an ROI with intracortical and subdural devices requires accurate assignments of position, orientation, and relevant limitations. For each solution of N devices, an array is generated containing three cartesian coordinates and associated rotational values describing the position and orientation, respectively. The six values per device remain in the MRI coordinate system units provided by the whole-cortex output from BrainSuite to maintain relation to sources. The combined array of devices and their positions is of array shape (N,6), which is the complete trajectory solution used in optimization methods.

With each newly generated solution, further refinement is performed with unique device specifications of RMS noise profiles, recording bandwidth, maximum trajectory angle, minimum and maximum depth in cortex, and proximity between devices, though additional options exist in the script. Limit corrections are looped until all are satisfied. Each device’s angle relative to scalp is approximated by the angle between the device trajectory vector and a center-of-brain to device vector (CD). The device trajectory vector is mixed with the CD until the angle between them is below the assigned limit. Depth for each device is calculated by reorienting the ROI vertices such that the z axis points toward CD and finding the farthest source in z. The distance between this farthest source and the CD point is an effective measure of maximum potential depth so long as only the cortex is considered. The device position is shifted along the CD vector to reach the desired limit in either minimum or maximum. Depth control is applied to assure that intracortical probes cannot be placed too deep and that subdural grids are maintained above the cortex rather than unrealistically within.

After all limit corrections are made and verified, source domain signal measurements are calculated as described previously. An array of voltages for each device relates to the maximum measurement possible for every source in the ROI, yielding null values outside of the lead field bounds to prevent calculation where not impactful. Each voltage array in source space is then converted to SNR and information capacity as described in §2.1, using individual noise profiles and bandwidth assigned for each device. The desired output value of voltage, SNR, or information capacity is then attained by taking the absolute maximum across all devices for each source point in the ROI. For optimization purposes, information capacity is summed for all sources in the ROI. For visualization, any of the three factors may be plotted on the ROI projected onto the ICS or PS.

Initialization of solutions is achieved with a customized gravitational repel method. Each device is randomly assigned a source location for its starting position. A cycle is then performed to allow devices within a defined proximity to repel each other with forces assigned by inverse square of separation, naturally spreading the distribution with variety on every repetition. Initial device angles are random, but trajectory limitations are still applied to create realistic implant trajectories. With this method, reasonable results are achieved without any optimization feedback by simply distributing devices throughout the ROI.

This format is usable with many optimization methods, but this study uses genetic optimization with the open-source Python library PyGAD (Gad, 2021). The shared code in §7 is provided with the settings used to achieve the presented results. In brief, genetic optimization settings used 30 generations, a population of 44 solutions per generation, the top 15 solutions are mixed for ‘mating’ where solution values are exchanged, mutation is performed varying by the inter-generational stability of the top solution, and limit corrections are made immediately after generating each population prior to scoring. To speed up processing, PyGAD’s built-in GPU parallel processing was used on a local desktop. The best results for each generation are saved along with the score for later plotting and visualization. Whole-brain and ROI visualization is enabled through a custom UI for entering device types and coordinates manually or by loading from an optimized results file. Surface reconstruction and visualization is performed with Open3D (Zhou et al., 2018).

### 2.4. Sparse sensor selection

Though information capacity is a valuable metric, it does not fully describe the diversity of information acquired from various devices in a source environment. It is valuable to describe ideal subsets of sensors with high diversity, as neighbors may contain redundant information. To search for more valuable arrangements, sparse sensing algorithms are utilized. In 2016, Brunton *et al*. demonstrated a method for sparse sensor placement by optimization of a classification task (Brunton et al., 2016). As demonstrated in Figure 1, this process involves a PCA applied to the dataset on the voltage axis, followed by LDA multi-class classification training to discriminate cortical column dipoles. In this application, each class is defined by a unique dipole or set of dipoles being active. The sensor space is then optimized for all available electrodes providing signal, specific to the provided device lead field(s) and combined arrangements (Figure 3). The result is a global classification accuracy for all input sources and a coefficient matrix relating each sensor to its contribution in classifying each source.

**Figure 3.**
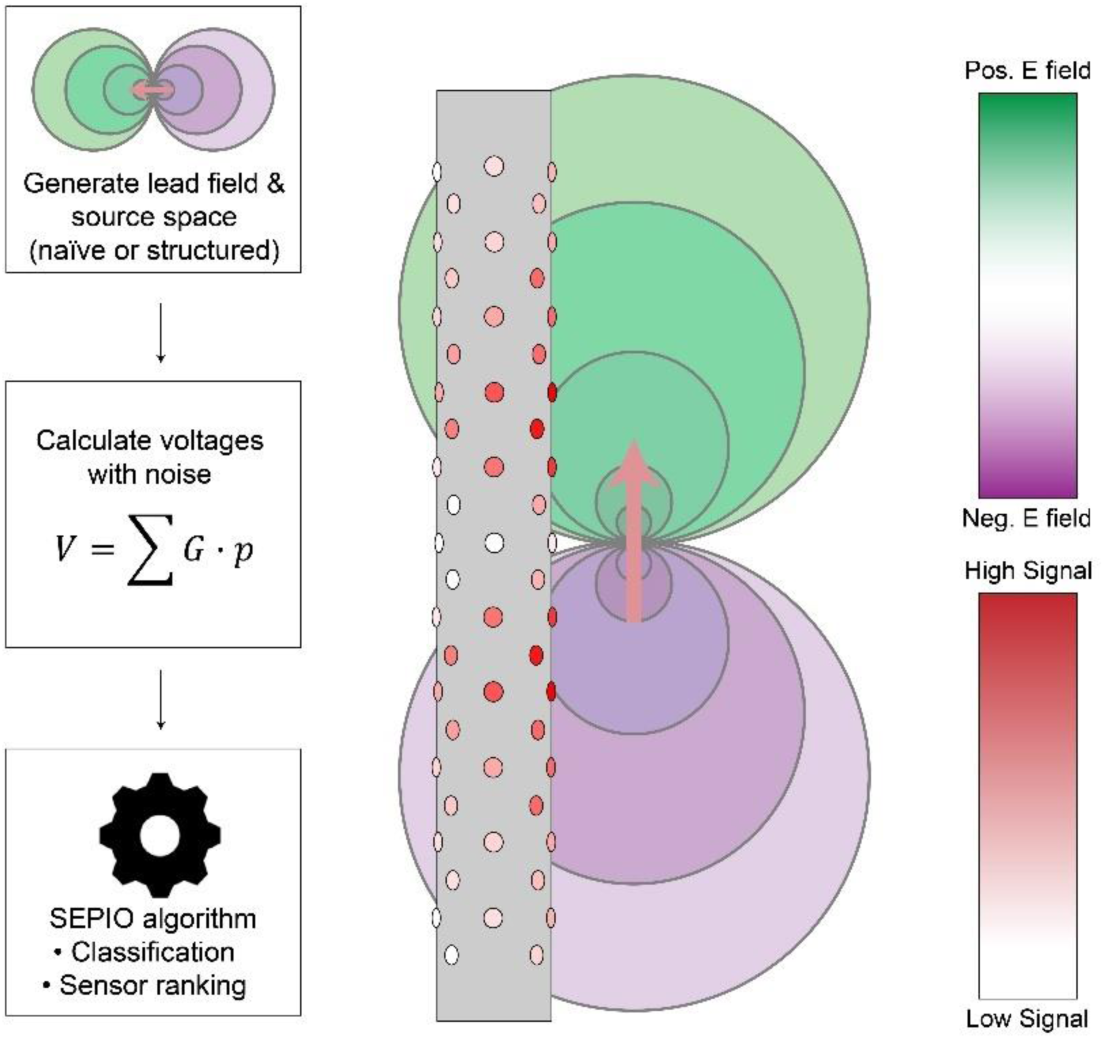
Method for simulated electrophysiology modeling with an example device and source. Lead fields are generated based on modelled devices and assumptions of surrounding conductivity. The source is modeled as current dipoles centered in cortex layer 4 with a standard current empirically validated. For each sensor, the simulated voltage is equal to the summation of dot products for all dipoles, *p*, and the lead field matrix *G*, demonstrated for a single dipole in the central image. The sensor array produces a voltage “image” that can be used for source classification and sensor ranking by SEPIO. Alternatively, the information capacity increases logarithmically with signal-to-noise ratio as measured by the maximum amplitude of a given device.

During the classification task, cross validation is performed to assess over- and under-training. Once training is successful, linear discriminant coefficients are used to sort the sensor IDs in order of classification priority (Brunton et al., 2016). This order is then used to select subsets for a second round of training. Top N subsets utilize the first sensors to retest the algorithm with the limitation that only those sensors exist. In this way, the highest quality arrangement can be compared to more limited subsets to evaluate the benefit derived from SEPIO. Visual assessment of the top subsets may also be valuable in determining inherent bias in the given system relative to signal strength bias.

Comparing simulated data from differing device types requires an additional step to balance the permitted compute power to the system complexity. Increasing the maximum sensors in a dataset increases the size of that dimension for both training and testing data, introducing a dimensionality challenge that only becomes apparent itself at larger values such as 200 sensors. To remedy this imbalance, simulated data is limited in the number of replicates per maximum sensors (RPS) such that some integer multiple of the dataset is replicated for both training and testing, provided unique noise generation for each to avoid data reuse. For example, in §3.4, this is performed with 1 RPS for DiSc and 8 RPS for virtual macros. With one device, DiSc trains 64 sensors with 64 replicates while virtual macros trains 4 sensors with 32 replicates. With five devices, DiSc trains 320 sensors with 320 replicates while virtual macros trains 20 sensors with 160 replicates. The higher training complexity with increased sensor counts necessitates higher replicate counts and prevents overtraining of smaller datasets. RPS selections were selected based on testing for adequate training without excessive replicate numbers for each individual device.

### 2.5. Source space generation with BrainSuite

Results presented in §3.1 are derived solely from manually defined source environments. In Figure 6, two tests are run in a Monte-Carlo fashion to evaluate effective cases for device choice and separation. In both of these tests, two SEEG or DiSc devices are simulated with randomly placed and oriented dipoles, and metrics are measured for information capacity in two manners. The best channel is used for each dipole to measure the maximum potential information capacity. For the first metric, values are recorded for individual device performance without intra-device montaging and averaged to find the mean single device performance. For the second metric, both intra-device and inter-device montaging are allowed to find the best possible channel pairs. The first test consists of the total dipole moment held constant, and the number of dipoles is increased. In the second test, the number and size of dipoles is held constant while the separation of two devices is changed in two different ROI sizes, 1 cm square and 10 cm square. Figure 7 explores the interaction of two devices when considering montaging effects by manual selection of channel pairs. Figure 8 continues with a similar idea applied to a mathematically defined curve of sources made to mimic a deep sulcus shape.

**Figure 4.**
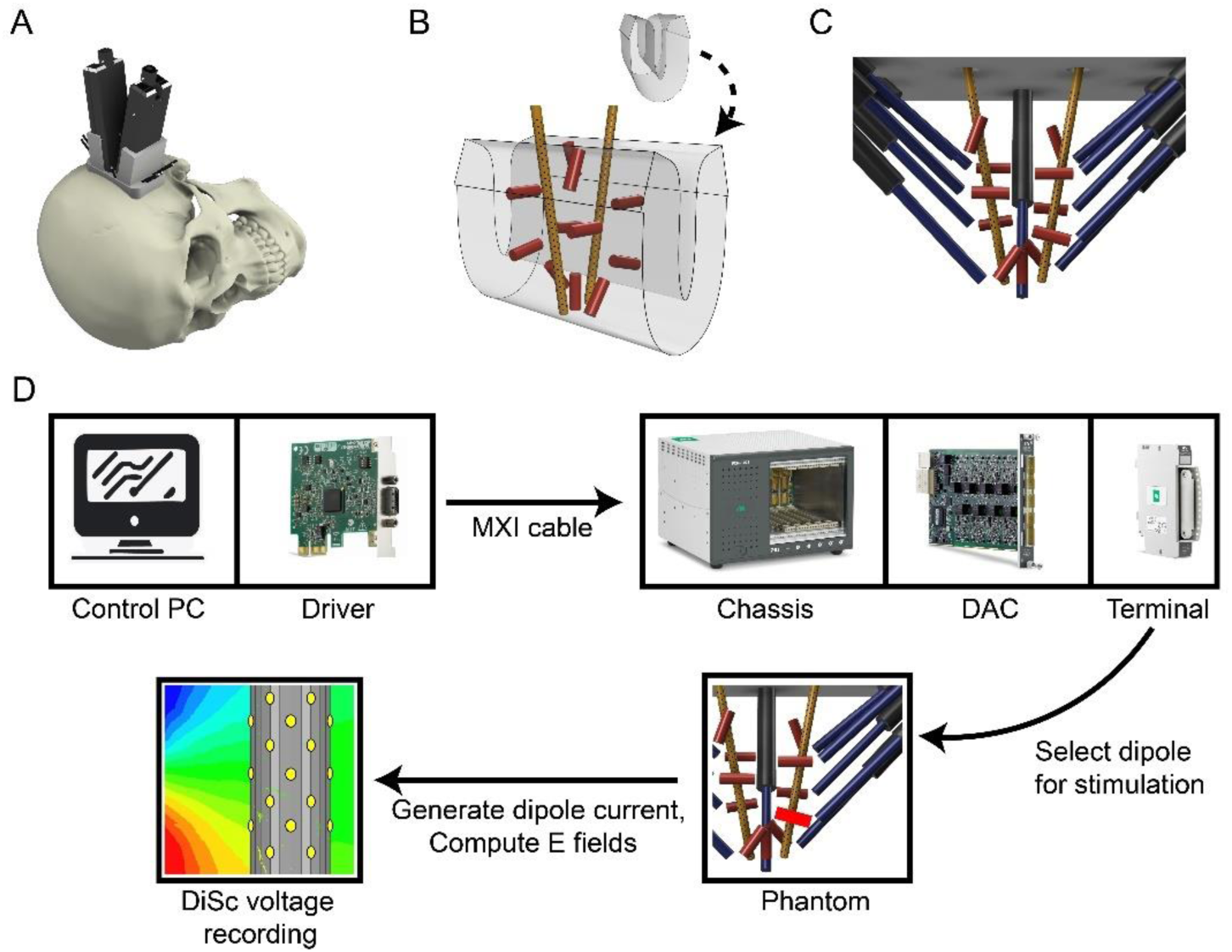
The design and operation of a phantom cortex model. First, (A) a head model is used to design a mount which supports all components including two DiSc devices in the same plane angled toward each other, as if placed within a section of the central sulcus cortex. (B) This cortex is used to place 11 (red) dipoles of 4 mm pole-to-pole separation, 10 of which are placed in planes ±3 𝑚𝑚 away parallel to the DiSc plane, and one of which is directly between the (yellow) DiSc probe tips. (C) (black) Model supports are then placed to guide (blue) wires into the proper location. Note that these dipole locations are simulated as intended and do not perfectly replicate the final construction. (D) The control PC runs NI-DAQmx which generates and communicates through the PCIe driver (NI PCIe-8363), MXI cable, and chassis (NI PXIe-1073) to instruct the operation of the DAC module (NI PXIe-4322). This module is interfaced by means of a terminal block (NI TB-4322), which is connected to 22-gauge twisted pair wire by screw terminal. This wire is connected as desired to a twisted wire pair with 4mm offset. During current stimulation, electric field generation induces a voltage on DiSc contacts and is recorded.

**Figure 5.**
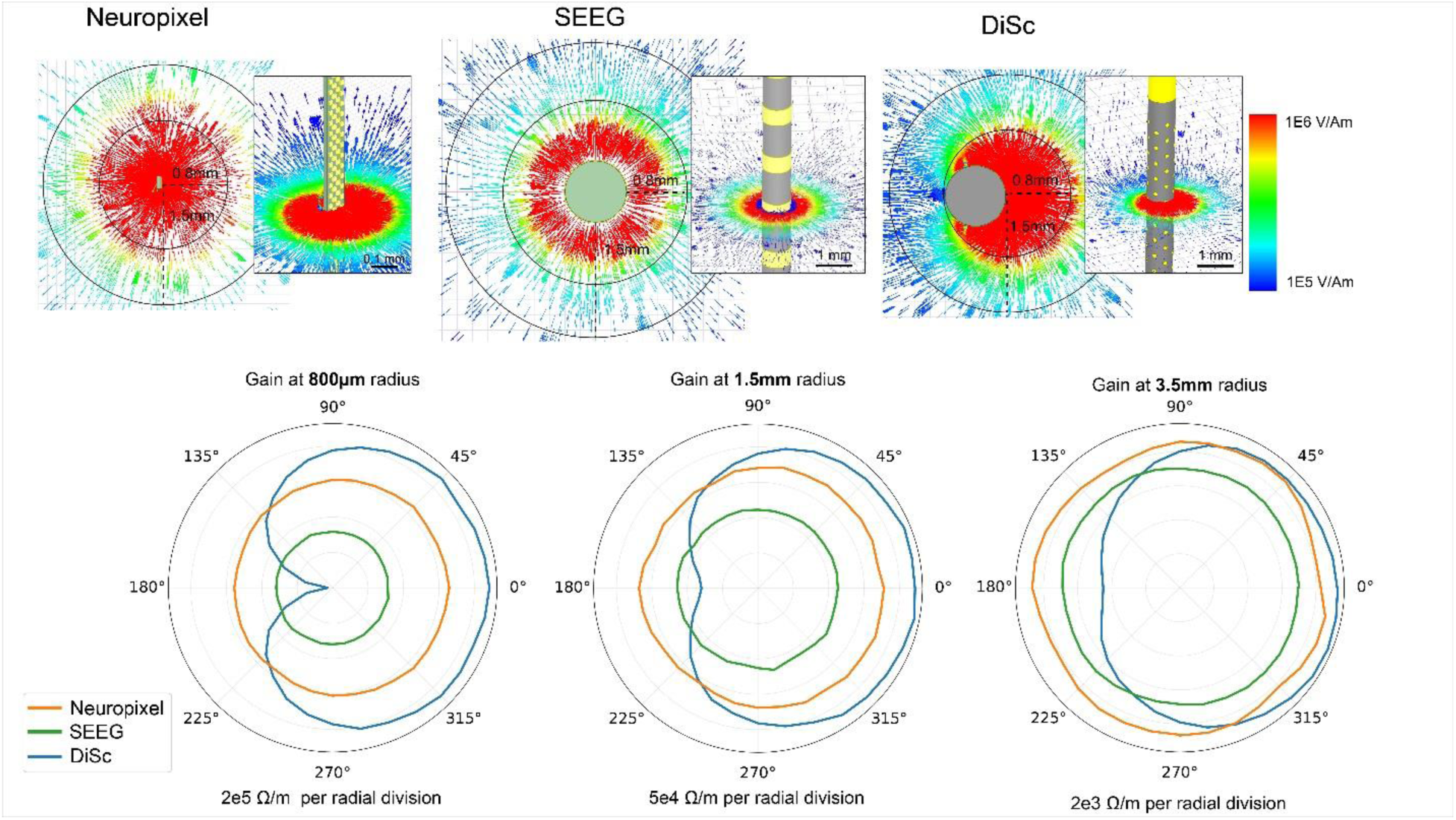
Demonstrated lead fields for three devices and the influence of the insulating body. (Top) Cross section of lead field sensitivity centered on a single contact for Neuropixel, SEEG, and DiSc displayed as a current vector field with the chosen contact centered in each display. (Bottom) Polar lead field sensitivity of a single contact at radii of 0.8 mm, 1.5 mm, and 3.5 mm for the three devices. Even at 0.8mm diameter, the micro-contact on an insulator can amplify sources nearby, while attenuating those on the opposite side. Radial resistivity scales vary per graph.

**Figure 6.**
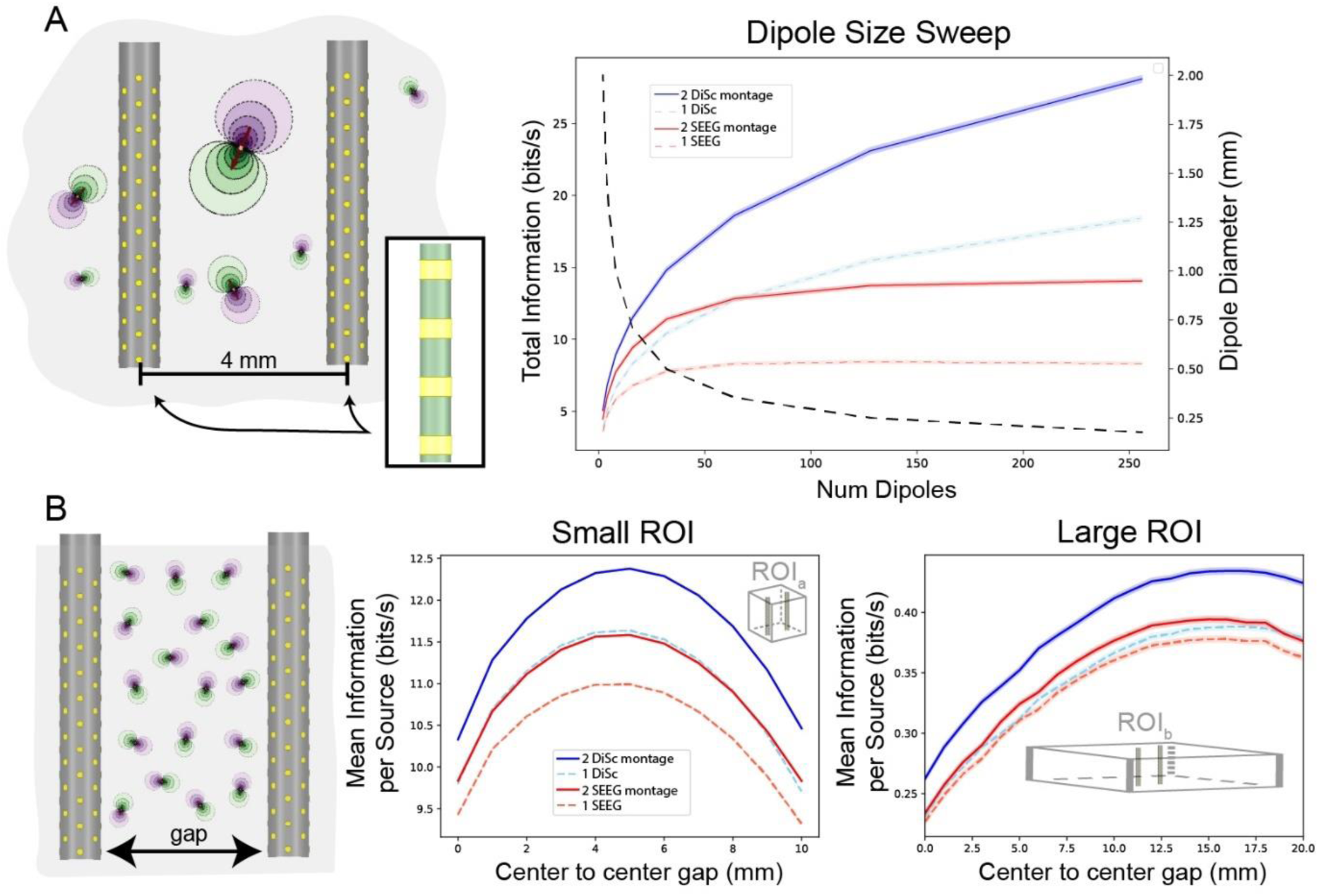
Information capacity from a generalized Monte Carlo simulation of SEEG and DiSc devices in a ROI with random dipole orientation. (A) The left image depicts the source space swept through the 1-256 dipoles while maintaining total dipole moment by changing dipole diameters with a constant device separation. On the right the dotted line describes the sources size and number on the right y-axis. In this environment, one and two devices of DiSc (128 channels each) and SEEG (18 channels each) are observed to determine the effect of bipolar montaging on total information measured in bits per second. (B) The left image describes a set number of current sources with stable magnitude and random orientation while device separation is changed. In a small ROI (1 cm square), peak information capacity is limited by the source space boundaries. In a larger ROI (10 cm square), the lower spatial density of sources lowers mean information per source but extends the efficient separation distance. In all graphs, the solid line surrounded by shaded region describes the mean and standard deviation derived from 10,000 Monte Carlo cycles.

**Figure 7.**
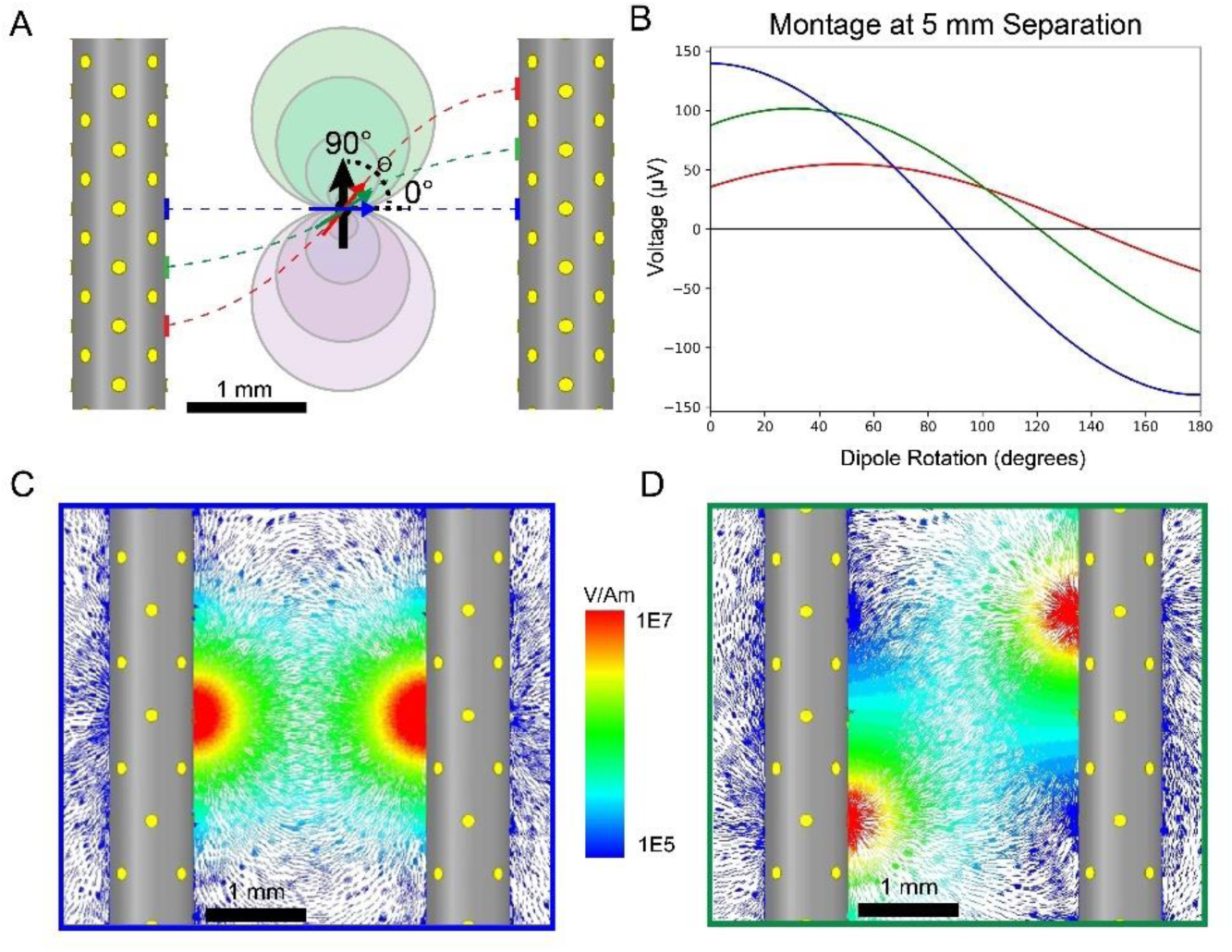
Montage quality is a function of contact pair sensitivity and source angle. (A) Three sensor pairs are chosen to demonstrate montaging between two devices 3 𝑚𝑚 apart with a central rotating dipole. (B) At different dipole angles, different montage pairs achieve the greatest power. Colors correspond to the pairs defined in A and peak power can be seen to decrease due to greater sensor separation. (C) Each montage pair can be observed in ANSYS simulation by placing a ±1 𝐴 current source on each contact. By reciprocity, the vector field (𝑉/𝐴𝑚) then relates any sources at those points to a voltage observed on the sensors. The blue montage pair is optimized for low angle dipoles while (D) the green montage pair is best for a rotation of around 40°.

**Figure 8.**
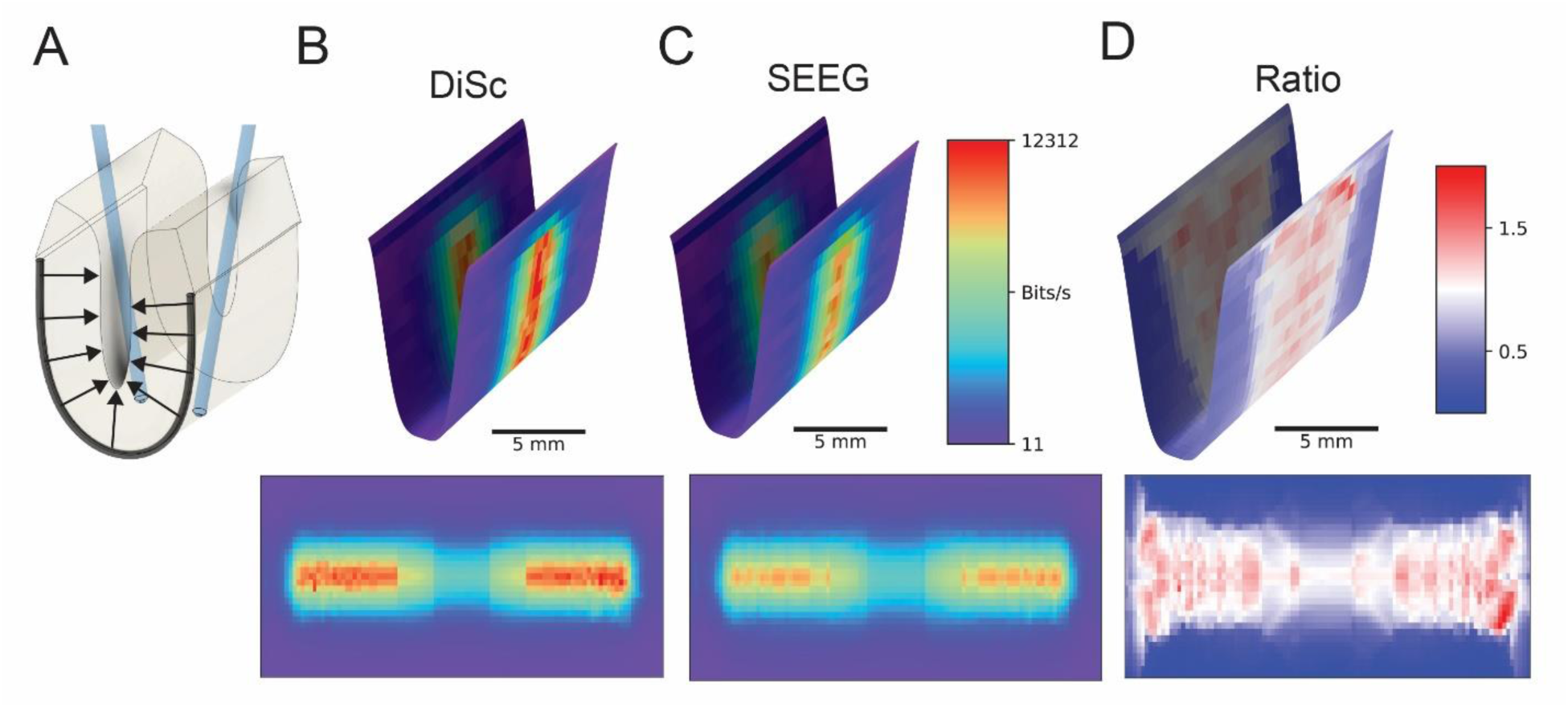
Information capacity projected onto the inner cortical surface for two different pairs of depth arrays in a central sulcus ROI. (A) Cortical column dipoles are distributed along an assumed sulcus as shown by arrows, with an SEEG-scale device on either side of the curve. Resulting values are projected onto the inner surface denoted at the front edge by a dark line. (B) DiSc and (C) SEEG devices are simulated passing through the source layer for each side of a modelled central sulcus where the signal is greatest. Results are shown in the (top) curved sulcus shape and (bottom) flattened cortical surface, with samples per second represented by the same color scale in both devices. (D) The point-wise ratio of DiSc to SEEG information capacity is shown to visualize their contrasting qualities.

BrainSuite was used to generate source environments throughout §3.2-3.4 using the scan 2523412.nii.gz from Beijing Normal University (CC-BY-NC) (Gao et al., 2022) under the International Neuroimaging Datasharing Initiative (INDI). Tutorial methods for cortical surface extraction are available on the BrainSuite website (BrainSuite, n.d.-a), providing details on recovering the ICS for subject-specific T1 MRIs. In order to extract only a desired ROI, surface and volume registration was used as described in the provided tutorials (BrainSuite, n.d.-b). Default values were used for registrations and extractions.

### 2.6. Phantom construction and SEPIO processing

Testing of SEPIO in §3.5 consists of the use of another manually defined source space in Figure 13 and a phantom model that is both built and simulated, presented in Figure 14. The phantom consists of a model head, a device fixture holding dipole wires and DiSc devices, a stimulation system, and a recording system. The design emulates a simplified section of deep sulcus, such as the central sulcus, with dipoles extending from shallow positions to 10 mm depth, offering a rich variety of locations and orientations.

The phantom is assembled using a PVC skull cut to accommodate the fixture housing the device and dipole wires (Figure 4). This custom fixture is designed in Fusion 360 and printed with a Form 3 in Form high-temperature resin (Formlabs RS-F2-HTAM-02). Twisted pair wires are supported by glass capillary tubes which are fed through the fixture supports, and wires bent to the desired site of dipoles along simulated sulcus. The wire is 30-gauge solid core, silver-coated copper and the final ∼10 mm extends from the capillary to reduce insulation effects and allow manipulation into the desired position. These wires terminate with exposed ends to form the two sides of a 4 mm dipole representing a simplified cortical column arrangement of deep sulcus. The ends of capillary tubes are sealed with a small amount of heat shrink tubing and silicone sealant. Next, two 64-channel DiSc devices are placed in the fixture. Images with a scale reference are then assessed with ImageJ to determine the arrangement of all dipoles (wire tips) relative to each DiSc, as well as DiSc rotation in order to recreate a matching simulation. Measurement variability is ±43𝜇𝑚 and rotation of devices is estimated to be within 15° of simulation, while simulation is limited to a 500𝜇𝑚 lead field voxel size. The skull is filled with a 0.15x PBS solution (Corning 21-040-CV) yielding a conductivity of approximately 0.25 𝑆/𝑚, representing grey matter (Koessler et al., 2017).

Phantom testing is performed with stimulation from a National Instruments system consisting of a NI PXIe-1073 chassis, NI PXIe-4322 stimulation module, NI TB-4322 terminal block, and NI PCIe-8361 for control by a local PC with NI-DAQmx software, as seen in Figure 4. One dipole is selected per trial and stimulated with bipolar current-driven sources with a magnitude of 5 𝜇𝐴 and a sinusoidal frequency of 20 Hz recorded for one minute each. This current through the local media sets up an electric field, generating a voltage at the recording device that can be used to classify environment states. This produces a dipole moment of 20 𝑛𝐴𝑚 which is larger than some baseline activity (Murakami & Okada, 2015), but is typical of evoked response amplitudes.

Data is generated by recording stimulation waveforms using DiSc for 1 minute at 20 𝑘𝑆/𝑠, downsampled to 2 𝑘𝑆/𝑠, and repeated for each individual dipole. DiSc recording is performed as described by Abrego *et al*. (Abrego et al., 2023) but utilizes an updated electrode and backend design. Data is pre-processed by phase shifting to a stimulation phase of zero at initialization and applying an order-5 Butterworth bandpass filter from 2 Hz to 118 Hz to remove DC and 120 Hz AC noise. The recording software also includes a built-in 59-61 Hz notch filter to remove electrical noise. Ambient noise is recorded for device assessment with the stimulation and recording equipment on and connected, but without providing current. Signal measurements with each dipole connected independently are used to assess the average RMS noise for this particular phantom arrangement. Overall, the characteristic RMS noise of DiSc was found to be near the prior 4.1 𝜇𝑉 value. This value is used to add similar Gaussian noise to the simulation dataset. Then, the simulation dataset is scaled by a ratio of the datasets RMS powers, such that their overall RMS power matches.

SEPIO trials are created from each dataset by capturing single timepoint signals at absolute power peaks for each stimulation period. A total of 1,208 trials per class are used for phantom datasets and 1,000 trials per class for simulation datasets. Trial arrays are randomly shuffled in correlation with their dipole class labels and supplied to the SEPIO algorithm with 80-20 or 60-40 train-test split used, depending on observed overtraining. Additional Gaussian noise is added to both datasets to increase training difficulty, yielding a final mean SNR of 2.7 for the phantom and 3.0 for the simulation datasets.

## 3. Results

We present device trajectory optimization in terms of total information capacity and source classification accuracy, which is part of the SEPIO tool. To achieve this, §3.1 provides a tool to assess and visualize the simulation assumptions. The source space and device lead fields are evaluated to observe the interaction between the two. In §3.2, we generate a ROI from a T1 MRI and project cortical dipole sources into the grey matter (Piastra et al., 2021). Optimization of multiple devices is then performed in §3.3 to determine the information capacity as compared to human trajectory selection. Following optimization, §3.4 applies SEPIO to the prior results to determine classification accuracy on over 3,000 cortical sources to validate signal coverage improvements and optimize sensor subset results. Finally, SEPIO is validated in §3.5 with simulation and phantom sources.

### 3.1. Information capacity, SNR, and device comparisons

The first step in simulation is deciding on devices that will be used to calculate voltage arrays. Each unique device is modelled and used to generate similarly unique lead fields. As such, it is best to compare a range of devices for each particular use case as their efficiency varies in detecting sources that are small or large, or oriented in a particular fashion relative to the sensor. Figure 5 demonstrates the clear differences in lead fields observable between devices.

Cross sectional sensitivity maps such as this aid in evaluating device design and directionality prior to simulations and without requiring physical construction. In this example, gain at 800 𝜇𝑚 displays an extreme directional sensitivity for DiSc due to the substrate shielding as seen in the top right of the figure. At further distances of 1.5 mm and 3.5 mm, the directional ratio is reduced but still present. Lead fields generated from AEDT are useful both in early device design guidance and are required for subsequent simulation for information capacity and placement optimization tasks.

We use the Shannon-Hartley theorem for information capacity (IC) as a natural metric for optimization. Lead fields and Eq. 2 allows us to compute the voltage amplitude at any contact due to any source. The contact size can also be assigned a noise floor inversely proportional to the surface area at a given bandwidth or frequency (Lewis et al., 2024). In this way, SNR for each source is used to compute IC using Eq. 1 and summed across all sources.

SEEG offers only axial variability while the directional lead fields achieved in DiSc offer a more robust opportunity for montaging but also presents common difficulties of microelectrodes such as higher noise profile. As such, the ideal distance between devices for information overlap is tested in arbitrary source space (Figure 6).

The dipole size sweep (Figure 6A) demonstrates the reliance on channel count and design in order to adequately sample more diverse source space. As the number of dipoles increases and the diameter shrinks, SEEG quickly approaches an asymptote as the larger ring electrodes are best at recording larger sources. Meanwhile, DiSc continues to gain total information capacity as sources become smaller, due to its smaller electrode surfaces. In considering the separation of two devices, it is important to consider the ROI scale used. A small ROI (Figure 6B) is important to validate that total information cannot grow further once devices are moved outside the bounds of the source space. In this example, it is easy to see a peak at 5 𝑚𝑚 where all devices begin to lose information capacity due to moving outside of the area that sources are generated. A large ROI (Figure 6B) can then be used to evaluate lead field montaging effectiveness in a more realistic scenario. Monopolar devices gain bits per channel up until the lead fields are sampling different spaces. Montaging two devices is similar at close distances due to oversampling but quickly gains value at distance. For DiSc, bipolar montaging values peak around 9 mm and then taper as further separation decreases the cross-sampling efficiency. Figure 7 further assesses the montage efficiency in terms of calculated voltage, relative to dipole angle.

Montages are computed by a subtraction of two sensor values for any given source. If one sensor lead field is subtracted from another, as visualized in C & D, any point in the space containing a source can be evaluated similarly as a montage of those two sensors. This comparison gives logical weight to the common use of SEEG linear serial montages but also enables more complex development for microelectrode arrays such as DiSc, particularly in classification, surgery planning, and source reconstruction. Though montaging requires two sensors for each measurement, single sensors can still be used to assess the general placement value based on SNR and the resulting information capacity. Figure 8 below demonstrates information capacity in a modelled central sulcus using intracortical DiSc and a virtual SEEG.

Spatial SNR and information capacity maps are a good way to visualize the sensitivity of recording devices in a known ROI, which provides an initial assessment of system performance for resource-intensive processes such as SEPIO. DiSc has a maximum information capacity of 12.3 kbits/s in the modelled central sulcus whereas SEEG has a maximum of 9.9 kbits/s. The total information capacity for DiSc is 7.71 Mbits/s and for SEEG is 7.53 Mbits/s, resulting in a difference of 179 kbits/s in favor of DiSc. Assessing the ratio map yields a point-wise comparison for greater detail. DiSc exceeds SEEG in the central region by up to 1.61x (890 bits/s DiSc vs 551 bits/s SEEG), while SEEG has an advantage of 0.47x for farther sources, but at a lower capacity magnitude (72.5 bits/s DiSc vs 154.2 bits/s SEEG).

While these maps highlight the differences in sensitivity of various types of sensors, it does not offer a comprehensive solution for the optimal placement of sensors due to the complexity of multi-device montaging and diverse tissue structures. To achieve accurate trajectory solutions requires accurate representation of tissue, which can only be achieved by subject-specific reconstruction of the source domain.

### 3.2. Subject-specific source simulation

T1-weighted MRI automatic segmentation is performed using BrainSuite as described in §2.2. In brief, a T1 scan is segmented to collect the inner cortical surface (ICSs), representing the surface interface between white matter and grey matter. Surface vertices and normal vectors are used to generate quasi-static dipoles, which reference device lead fields to generate a simulated voltage.

A global assessment of the exported ICS is performed in Figure 9 by mapping the relative angle difference between the ICS normal and the nearest scalp normal in a point-wise fashion.

**Figure 9.**
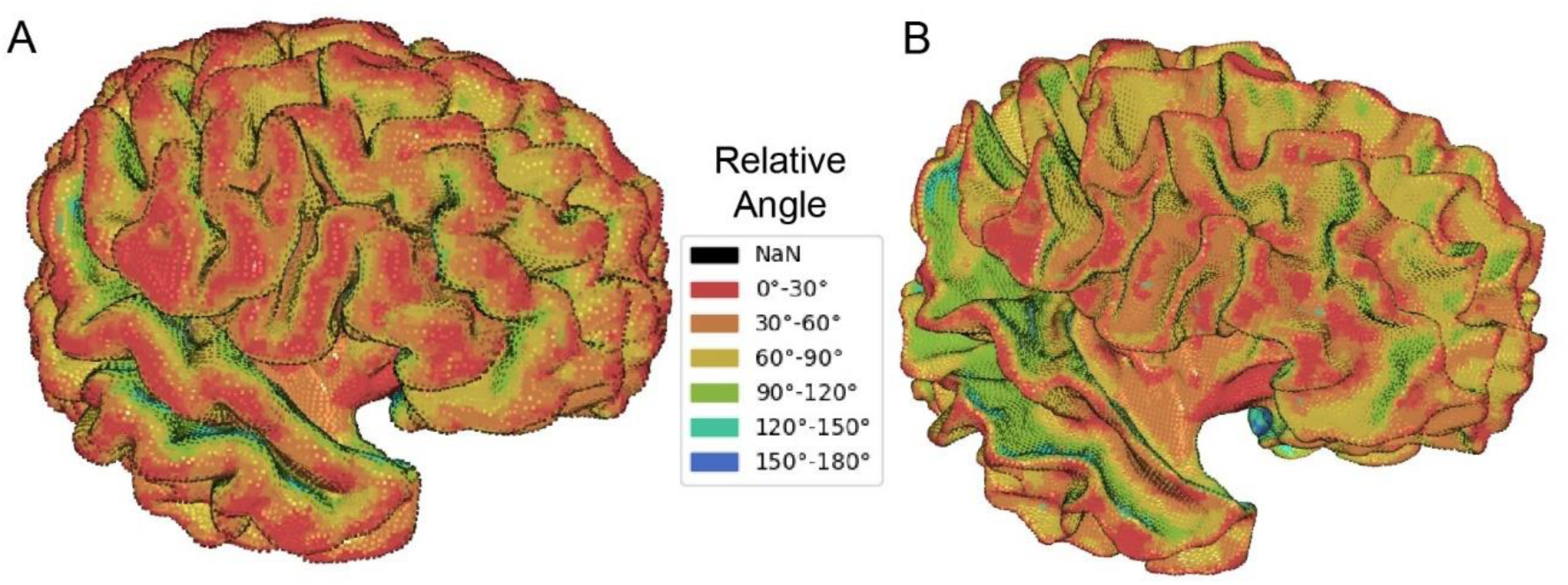
Heatmap of normal vector angles relative to scalp on reconstructed cortical surfaces. Reconstructed MRI surfaces are shown with (A) inner cortical surface values projected to the pial surface and (B) the inner cortical surface itself, allowing better observation of deep sulcus structures. The heatmap shown is applied pointwise to each vertex by assessing the difference between its normal vector and the scalp vector as described in §2.2.

Scalp-relative angle heatmap provides little benefit for intracortical devices but does have potential application in EEG and MEG. Sources parallel and anti-parallel to the surface are sensitive to EEG while orthogonal sources are sensitive to MEG. However, this provides the benefit of demonstrating the pial surface (PS) and ICS. The PS is a more common depiction of the cortex, but the ICS provides benefit in viewing highly contoured areas of sulcus. This demonstration also helps visualize how device lead fields will measure the sources represented by each point and normal vector. Each device position is used to shift the sources into its reference frame and calculate voltages (Eq. 2) in source domain or sensor domain, depending on the application. Figure 10 visualizes a single device placement with a source domain heatmap.

**Figure 10.**
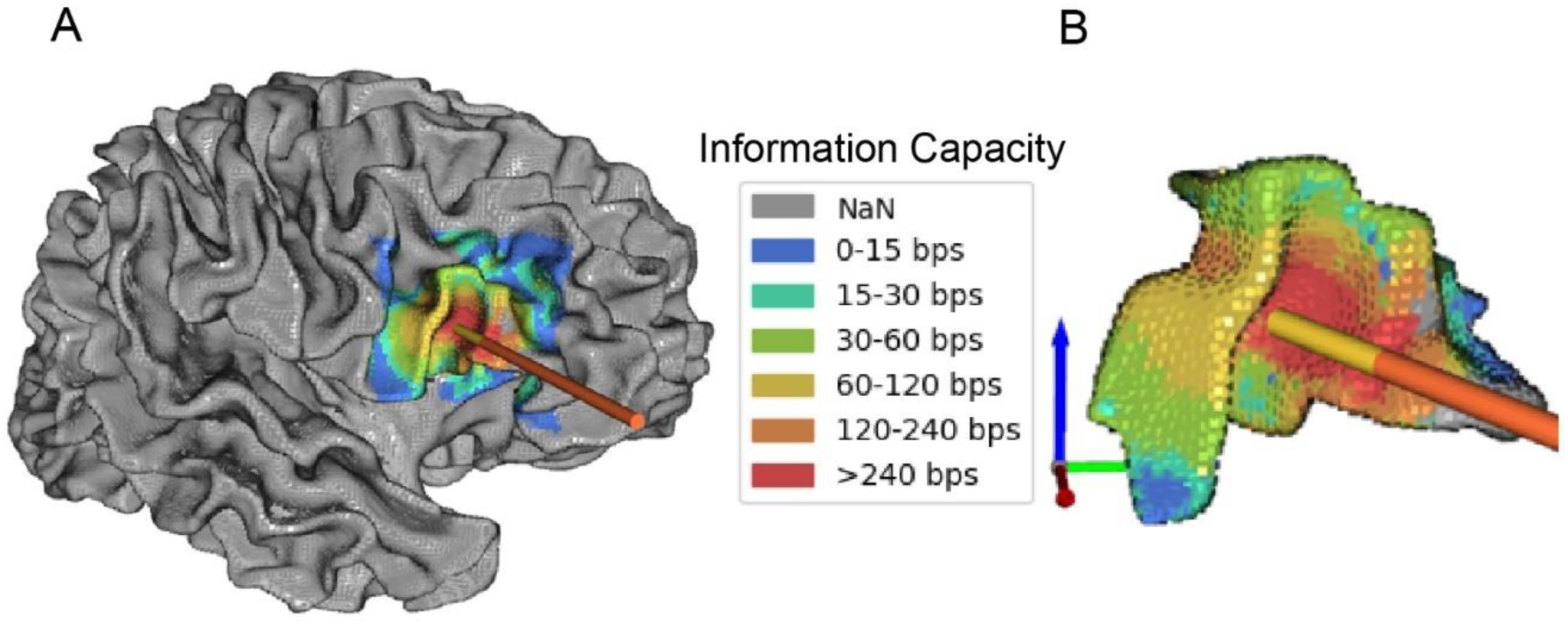
Heatmap of source domain results demonstrated with one DiSc. The (A) whole brain and a (B) specific ROI are mapped pointwise with the maximum potential information capacity (IC) for the device arrangement provided. Here, DiSc values are calculated with RMS noise of 4.1 µV and a bandwidth of 100 samples per second.

In this manner, the source domain can be viewed for any number of devices of any type by measures of voltage, signal-to-noise ratio, or information capacity. Without any further processing, this enables a manual estimation of sites with good sampling and sites requiring further improvement. The same can also be performed by plotting source domain statistics and comparing solutions for any ROI or the whole brain. For optimization purposes, a single metric is derived from the information capacity by taking the sum over only the ROI sources. Since the total information capacity is determined by the pointwise maximum independent of devices, and because it is reliant on device characteristics, this unit of measurement can adequately compare trajectory solutions.

### 3.3. Multi-device trajectory optimization

Using the simulation system described prior, feedback can be achieved to improve results based on a variety of desired factors. Since simulation is device agnostic, it is important to consider frontend and backend limitations in constructing a fitness function. In this study, information capacity is used as the fitness metric since it additionally contains information regarding the recording system of each simulated device. To briefly summarize the optimization process (§2.3), each solution contains the position and orientation of every desired device of any type and number relative to the ROI. To initialize this process, random positions are chosen, and a simple algorithm repels device positions to distribute coverage. This method is tested against human selections. The best of 100 initialized solutions beat the best of five human solutions by 18.5% total information capacity in the same ROI shown below. Considering the computational requirements and stochastic nature of these methods, it is still suggested to provide initial seed parameters from human selections. In this manner, the remainder of the population can be filled with the best stochastic results and be optimized with intelligent solutions as a catalyst. Solutions become drastically more convoluted with increasing device count and type such that human aid will likely always provide benefit in reducing initial computational load. With the initial population of solutions generated, further optimization is performed using a genetic algorithm and results are shown for 1 to 5 DiSc devices in a Broca’s area ROI in Figure 11.

**Figure 11.**
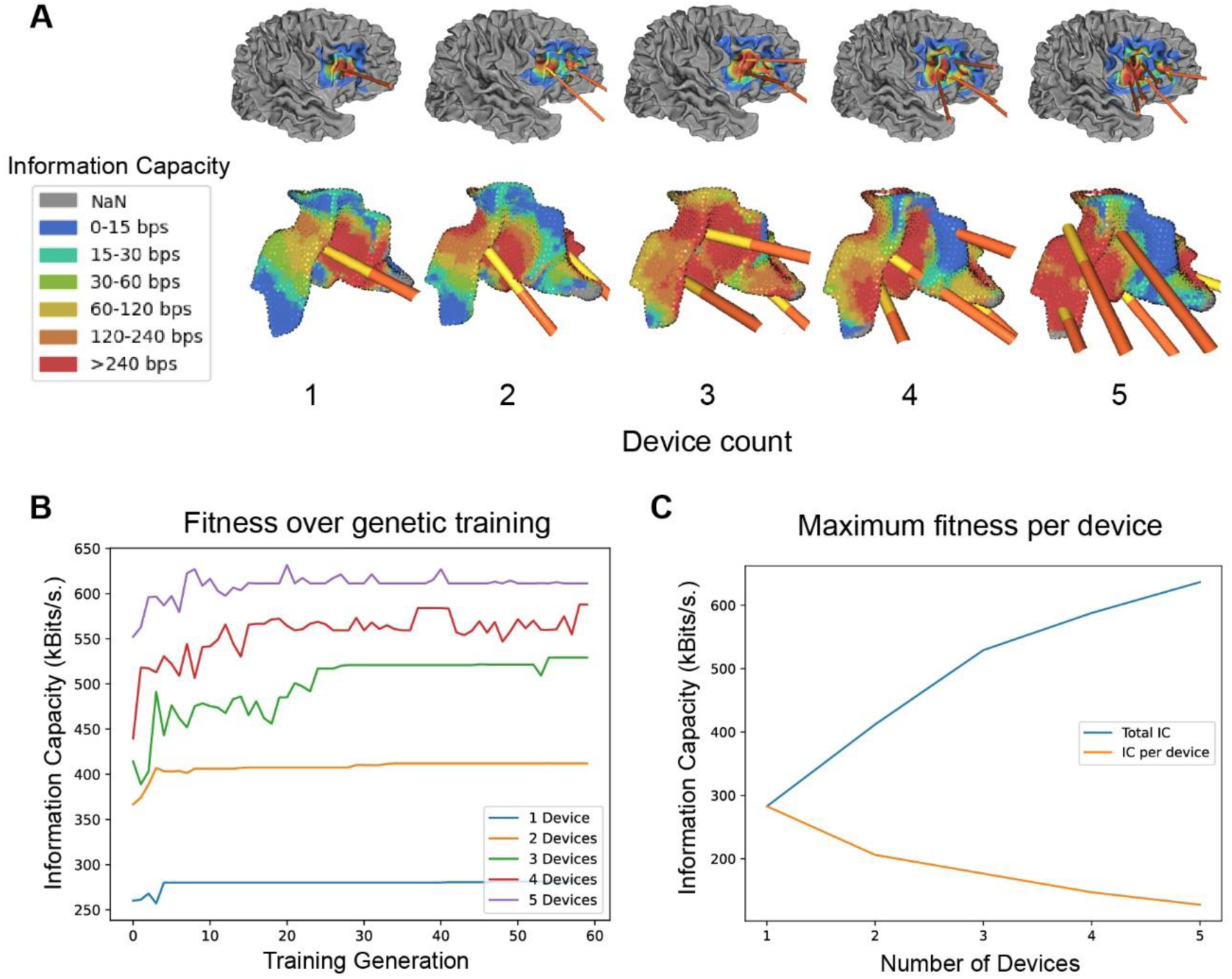
Genetic optimization of 1 to 5 DiSc devices in a right Broca’s area ROI. (A) The final solutions for each optimization are show on isolated cortex with heatmap showing the point-wise maximum voltage. (B) Information capacity as a fitness score is shown for each device count with maximum score plotted at each generation. (C) The maximum fitness observed over training is plotted in blue for each device count and the fitness gained per device is shown in orange. Signal montaging is not performed in this experiment and device trajectories are limited within 45° relative to the scalp normal.

Total information capacity increases with device count in a near-linear fashion likely since the majority of signal is derived from sources within ∼2 mm and devices are spaced at least that far apart. However, there is a notable diminishing return in the total information capacity per device since lead fields overlap and will gain no further benefit from repeat sampling at the same sensitivity without montaging. An additional consideration for increased device count is computational expense. Genetic optimization is performed in parallel on a GPU and required five hours for the entire process of one through five devices, with processing time increasing superlinearly with each added device. However, this can be mitigated by strong initial populations, such as those defined by expert human input, which begin the process with high fitness and use optimization only for fine tuning. Once a terminal solution is achieved, visualization is a good human-readable quality assessment for coverage. To further assess the solution, classification methods are employed.

### 3.4. Sparse sensor optimization and classification of trajectory results

Optimized solutions for each device group have been tested in the source domain above and will be evaluated further in the device domain. Sensor voltage arrays are reproduced for each source sequentially with labels generated for training and classification. SEPIO is then applied to classify and provide further optimization information on each trajectory set. This process provides both the observed classification accuracy for each source which is averaged in Figure 12 for each number of devices per solution and sensors permitted to classify.

**Figure 12.**
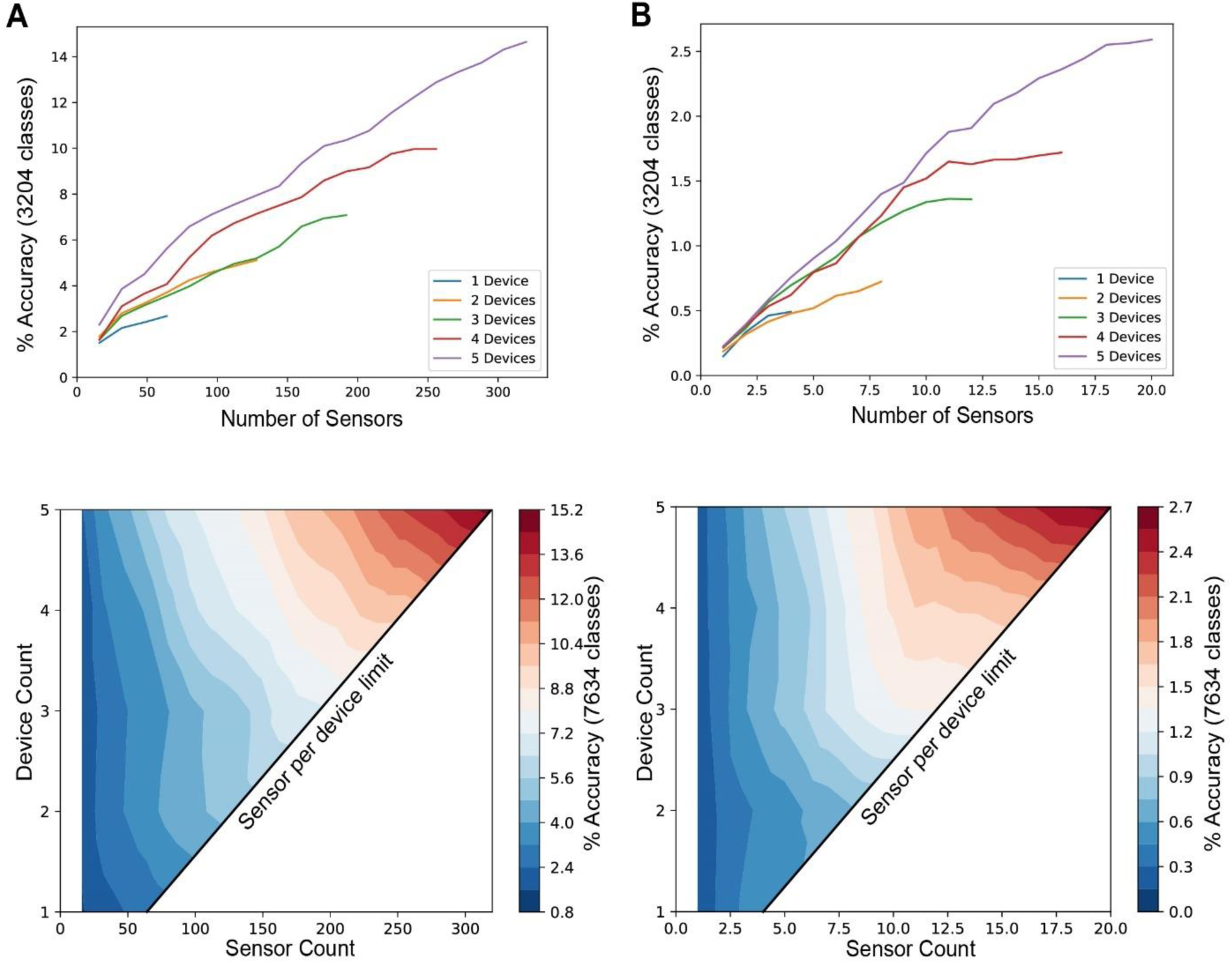
SEPIO applied to optimized trajectories of 1 to 5 devices in Broca’s area containing 3,204 sources/classes. Top trajectory results for each device count in Figure 11 are used as an input for SEPIO using either (A) DiSc devices or (B) virtual macro contacts calculated from DiSc devices for fair comparison. (Top) Each trajectory solution is represented as its own line with sensor count increasing as sorted by SEPIO training. The classification accuracy of the test set at each point of the line is used to create a (Bottom) heatmap of accuracy based on the allowed device and sensor count. Sensor count is incremented by 16 and 1, respectively, and heatmap contour lines are blended between the defined points.

**Figure 13.**
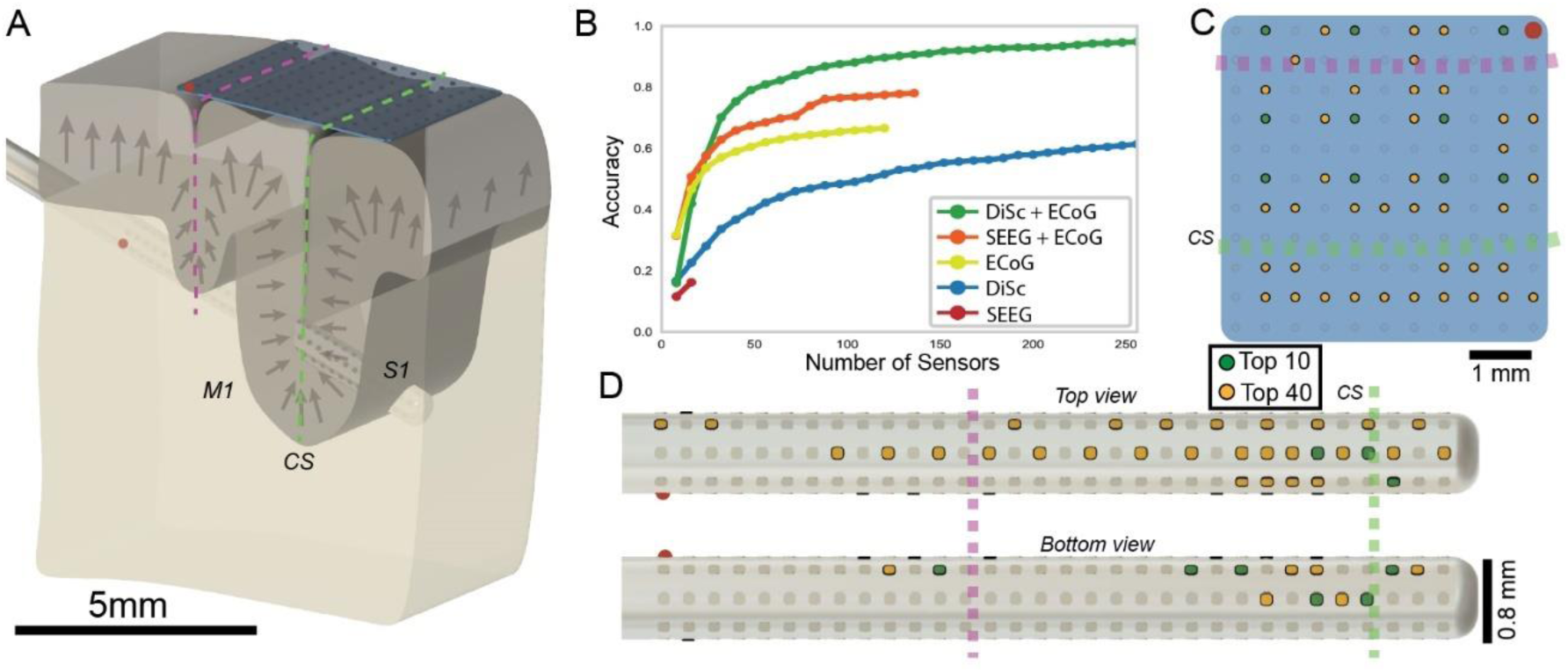
SEPIO performed on a simulated two sulcus model with various recording devices. (A) The two-sulcus model is constructed with 128 dipoles, activated independently for the classification task. (B) Devices are tested individually and in combination to assess the SEPIO classification performance. Data points are only shown on every 8 sensors. (C&D) For the best performing combination, the top ten sensors per device are shown in green and the top remainder making up the top 50 are shown in yellow. An orienting red dot is shown on the (A) model devices, (C) ECoG grid, and (D) DiSc. Dotted lines in yellow and green are also used to mark the locations crossing sulci.

**Figure 14.**
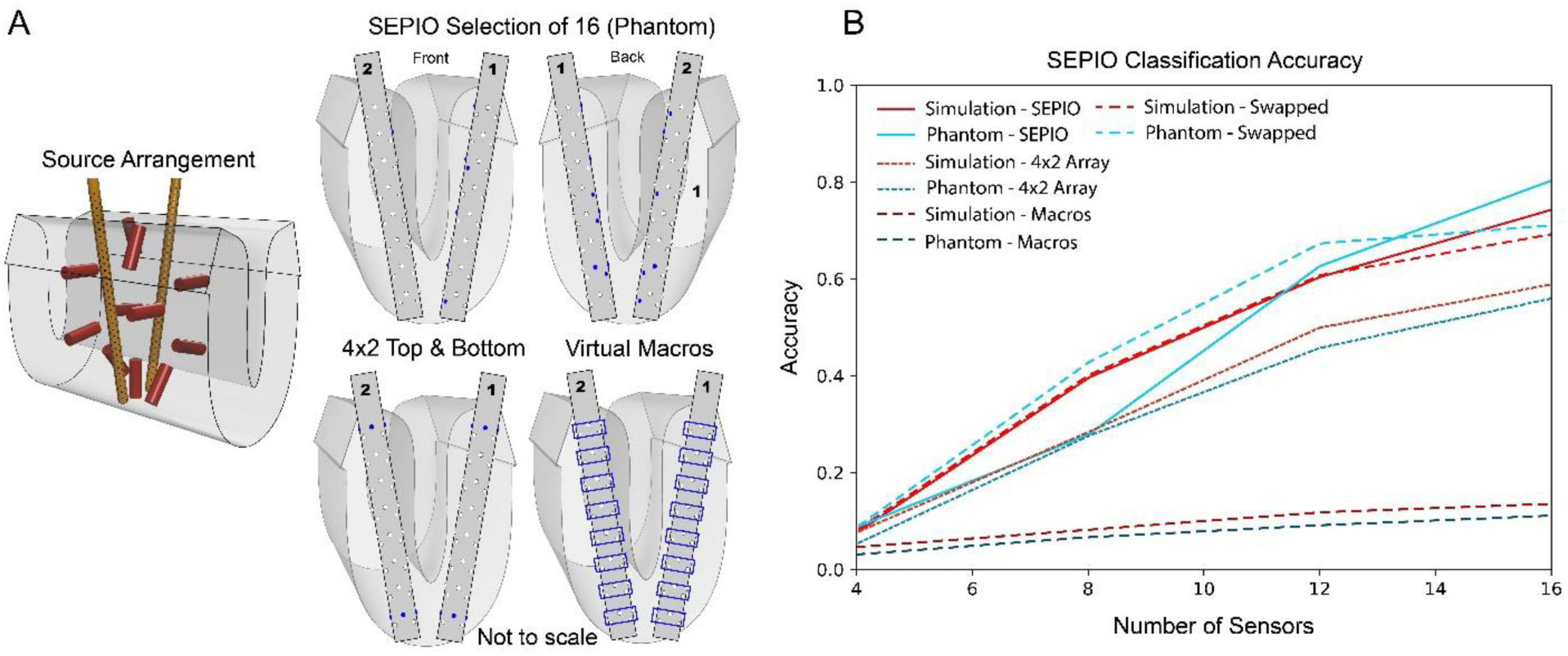
Classification accuracy of the phantom arrangement reduced to 16 sensors by SEPIO versus human-defined subsets. Sensor configurations are visualized for each case. The DiSc subset displays both sides of the SEPIO selected arrangement; (bottom row) 4×2 pattern and an SEEG 1×8 show only one side. (B) SEPIO classification is applied to the whole sensor set and two pre-defined subsets of 16 sensors. Phantom and simulation top 16 limited sensor lists are also swapped for validation. Training is performed as before except for realistic noise reduction for virtual macros. With 55 classes, chance is 1.8% accuracy.

Assessment by SEPIO shows that increasing device count, and thus spatial diversity, allows the same number of sensors to provide more information. For DiSc (Figure 12A), an imagined vertical line at 128 sensors measures around 4% accuracy for 2 devices and 8% accuracy for 5 devices. Allowing the maximum sensors for 5 devices exceeds 15% accuracy on 3,204 source classes. From the standard DiSc array, virtual macros are computed to represent a similar format to a 4 channel SEEG by averaging voltage in rings of 16 microelectrodes. An equivalent vertical line at 8 sensors measures about 0.6% accuracy for two devices and 1.4% for five devices. Allowing all sensors reaches an accuracy of 2.7%. This large difference in scale is also in relation to the sensor count, as the low sensor numbers for DiSc are comparable. Looking at 16 sensors, DiSc achieves a maximum of 2.3% accuracy compared to 2.4% in virtual macros.

The sorting of individual sensors is useful for custom device design and further manual solution refinement. For example, DiSc devices hold 128 contacts on the probe body but can typically only record 64, allowing the option for customized probe designs similar to NeuroPixel. When multiple device types are employed, this provides a measure of effectiveness for each, aiding the decision for the number of invasive vs non-invasive devices, trajectories, and their types. Subset sorting may also improve live recording visualization by emphasizing channels with the best expected signal.

### 3.5. SEPIO multi-device demonstration

The performance of SEPIO in simulated T1-MRI is successful, but must be validated and compared with a simplified model. A simulation is used to evaluate functionality in handling multiple device types and a real-world phantom of 11 sources is generated and compared to a matching simulation. SEPIO is first assessed in a simulated two-sulcus model with a variety of recording devices. 128 dipoles are arranged across four rows, oriented normal to the grey-white interface and activated one at a time for classification. Figure 13 visualizes two intracortical devices, DiSc and SEEG, and one subdural ECoG grid, used in various combinations to explore individual and synergistic sensor value.

Individual devices performed relative to their sensor quality and source proximity. SEEG performed poorly below 20% accuracy, in part due to the limitation to 16 sensors. DiSc performs better, reaching 57% accuracy at 256 sensors. ECoG gets the most value from the first 48 sensors passing 60% accuracy and later 66% with all 128 sensors. The combination of these devices allows better spatial coverage of the space and permits SEPIO to choose from either device in any order to optimize the classification. Utilizing SEEG and ECoG meets 76% with all 146 combined sensors while DiSc and ECoG reach 89% at the same point, and 93% at 256 sensors. This simulation approach demonstrates the practicality of sparse sensing for LFP classification, as well as the value of multiple device arrangements to diversify lead fields. However, there is still a necessary validation to be made with real-world data as compared to simulation.

### 3.6. SEPIO validation with simulation and phantom

To make a comparison to human-designed models, a SEPIO selection of 16 sensors is compared to simplified arrays on the two DiSc. This effectively compares sparse optimization of the dense grid to naïve sparse sensor selection. The 11 dipoles are used in a choose-2 manner, generating 55 distinct classes. In this test, SEPIO is allowed to sample all 128 channels to choose a top 16 used for classification comparison. This is compared to SEPIO classification permitted to train and test only a simple array of 16 sensors and a ring electrode model where rows of contacts are averaged into virtual macroelectrodes (Abrego et al., 2023; Sun et al., 2022). Figure 14 demonstrates the classification accuracy and sensor distributions for SEPIO versus regular contact configurations.

SEPIO is successful at selecting a sensor subset, achieving 75% and 83% accuracy for simulation and phantom, respectively. In comparison, the 4×2 array reached only 59% and 55% accuracy, while virtual macros suffer from signal averaging and yielded 14% and 11% accuracy, respectively. The success of the 4×2 array is a function of the 11-dipole arrangement but demonstrates the effectiveness of small contacts on a sufficiently large insulating body so as to provide diverse signal inputs (Abrego et al., 2023). This distribution is reasonably successful but is 22% less accurate than the SEPIO subset on average. Though the virtual macro array is given a lowered noise profile, signal averaging impaired results. The macros have similar axial diversity as DiSc but performed poorly. A U-probe configuration (Plexon Inc.) using DiSc microelectrodes was tested as well (not shown) and resulted in 65% and 73% accuracy on average, which is also less than SEPIO-selected microcontacts.

The two trials marked ‘swapped’ involve the training on full datasets (128 channels) and the reduction of the test sensor sets to the top 16 selected by the opposing process. In this way, the top 16 selected for simulation are applied to phantom testing and vice versa. Simulation swapping lost 5% accuracy and phantom lost 9% accuracy. This is within reasonable limits given the difficulty of *in silico* reproduction of physical sources and contact angles. Some points during the increasing sensor count, swapped accuracy exceeded the full dataset, an interesting effect discussed further in §4.3.

Note that too many sensors selected for sparse methods may lead to oversampling and high accuracy, but in that situation any contact arrangement will be reasonably successful. Supplemental Figure 1 and Supplemental Table 1 further explore this idea using 32, 24, and 16 channels with a wider variety of defined arrays. The number of sensors appropriate for a ROI is also related to the diameter of the assumed source.

## 4. Discussion

The results of this study demonstrate the effectiveness of trajectory optimization and SEPIO in processing biophysical simulation data to enhance device arrangements and identify optimal sparse sensor subsets. Our method offers a complementary approach to the current implantation strategy, which is to avoid critical vasculature and place contacts in the region of interest (ROI). By improving the analytic value of local field potential (FP) recordings, within subject-specific ROIs, using realistic devices, we aim to elevate the quality and diagnostic potential of electrophysiological data in clinical settings.

### 4.1. In silico voltage imaging utility and limitations

Information theory has long been applied to single-unit neural activity but not to the optimization of recording array design (Bartolo et al., 2020; Shannon, 1948). While placing contacts for point-like single units is straightforward, optimization for high counts of spatially extended sources like multi-unit activity or local field potentials (LFPs) is non-trivial. Using LFPs as a proof-of-concept, we argue that the choice between individual measures, e.g., LFPs versus single units, should be reframed as a mission to maximize the total information capacity of recording arrays across all available biophysical sources. Achieving this goal requires models that incorporate the spatial extent of the target source and account for redundancy arising from mutual information between channels. The key biophysical parameters for such a model are summarized in Table 1, with relevant source scales and frequency bands illustrated in Figure 15.

**Figure 15.**
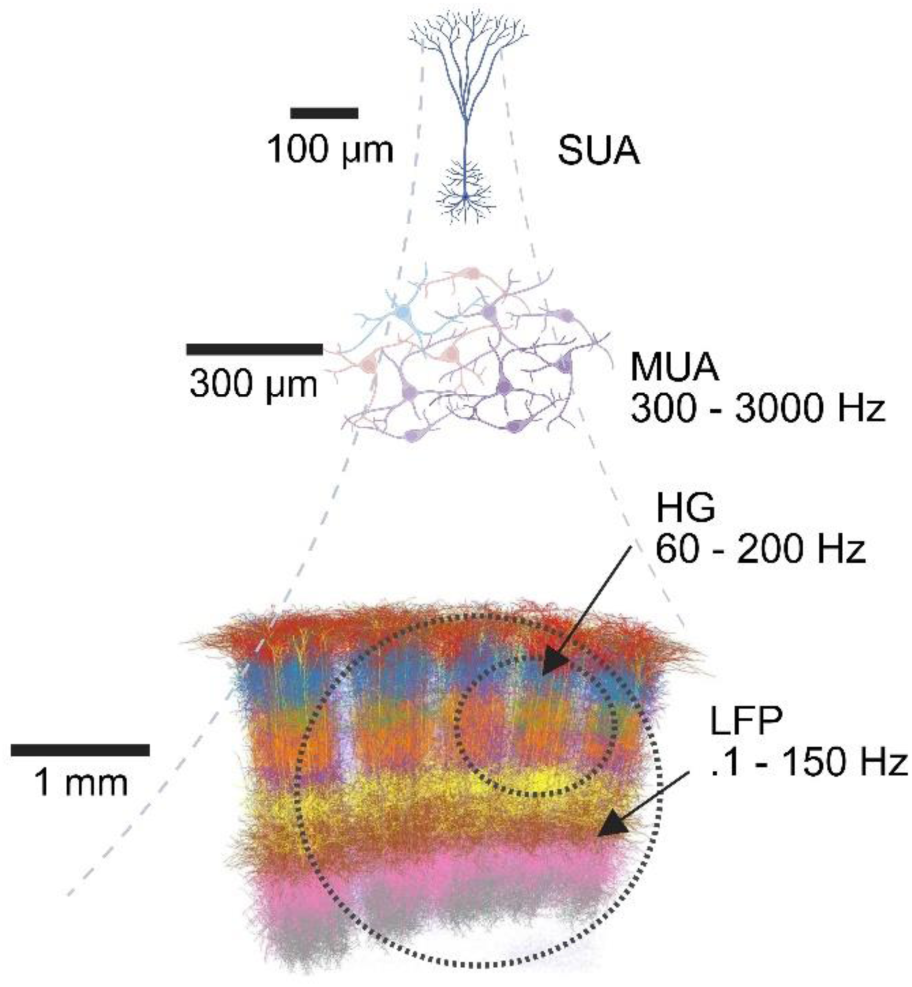
Neural sources displayed with relevant spatial scales and source bands. Theoretical Shannon-Hartley limit is calculated using the effective coding bandwidth and the high end of typical SNR, with a conversion from 10log10 to signal power ratio. This figure was made with BioRender.com and an image released with CC BY-SA 4.0 in 2022 from Marcel Oberlaender.

**Table 1.** Towards a useful model of multi-modal cortical information sources. Values are estimated only for cortical sources using the cited sources discussed below. Effective coding bandwidth selections are discussed in the following paragraph.

| Biophysical Event | Effective Coding Bandwidth (Hz) | Typical SNR (V/V power) | Theoretical Shannon-Hartley limit (calculated bits/s) |
| --- | --- | --- | --- |
| LFP (evoked potentials) | 50* | 8-20 | 220 |
| High gamma | 100* | 2.5-5.0 | 258 |
| MUA | 400* | 1.8-3.2 | 828 |

Values marked by an asterisk (*) are our estimates for cortical sources, as these metrics lack broadly accepted standards. Unlike the agreed-upon filter bandwidth, the *effective coding bandwidth* approximates the rate of actual usable information, where signal power is typically concentrated by the 1/f power law and biophysical encoding limits. For LFPs, this bandwidth is informed by studies of maximum evoked responses, such as visual evoked potentials (Kothari et al., 2016), which occur from 10 to 100 Hz with strongest responses in the mid-range. Based on this evidence, we adopt 50 Hz as the effective coding bandwidth. High gamma presents a more nuanced issue as the source and range (Jia et al., 2011a; Leszczyński et al., 2020) are more varied, though it is expected to be below MUA with reported power around 100 Hz (Lundqvist et al., 2016). For MUA, the bandwidth of 400 Hz can be explained as the fastest firing rate of the major signal sources, such as pyramidal cells and fast-spiking interneurons with a sustained rate of 200 Hz with bursts of up to 400 Hz (Kawaguchi, 2001). These assumptions are based on limited data points to be further investigated, and are ultimately unique to the experimental data. As such, the effective coding bandwidth can be assumed for a mid-range value, but is ultimately defined strictly by experimental data unique to each recording environment and subject.

Spatial extent of biophysical signals is critical for predicting mutual information in future versions. LFP spatial extent is in the centimeter plus range (Kajikawa & Schroeder, 2011). High gamma generators are generally assumed to be on the order of 300 µm but can generate activity that is volume conducted up to several millimeters from the source (Gieselmann & Thiele, 2008; Jia et al., 2011b). MUA sources are known to be local (Kajikawa & Schroeder, 2011; Mineault et al., 2013) and some estimate a volume conduction range of up to 200 µm (Arezzo et al., 1981; Harris et al., 2016). Typical SNR is the last component to calculate the information capacity limits. For LFPs, *in vitro* and *in vivo* experiments have determined an expected SNR of 9 to 13 dB (8-20 V/V power) (Abrego et al., 2023; Suarez-Perez et al., 2018). High gamma sources in the motor cortex have been observed between 4 and almost 7 dB (2.5-5.0 V/V power) (Kacker et al., 2025). MUA sources have been observed around 2.5-5 dB (1.8-3.2 V/V power) in murine cortex (Fiáth et al., 2016), though it has been suggested that values below 4 may not be consistent (Stratton et al., 2012). These parameters were applied to the Shannon-Hartley theorem (Equation 1) to compare the relative limits, but further study is needed to refine these estimations. For example, the theoretical information capacity for typical LFP sources is given by the effective coding bandwidth (50 Hz) multiplied by the base-2 log of the maximum SNR in V/V power plus one (21), yielding 220 bits/s.

The results observed in §3.1 demonstrate the information capacity with varying dipole size, inter-device distance, and dipole angle using two devices either individually or montaged. For both modeled intracranial arrays, capacity increased as the number of dipoles rose and their diameter decreased. DiSc showed a greater increase and continued rising, whereas SEEG plateaued at approximately 128 sources. Increased separation of two parallel devices observed a definitive increase in information capacity (Figure 6) up to a point of best coverage determined by comparing the small and large ROI. In all cases, performance is improved when considering montaging of sensors rather than in monopolar (distant reference) mode. Consequently, for any given recording environment there exists an ideal positioning for montaged sensor arrays. This optimal placement maximizes information capacity by balancing synergistic detection at the device pair’s center with diverse detection of the surrounding volume. Therefore, known cortical architecture can be used to define ideal device spacing and angles, providing refined inputs for machine learning optimization of sparse sensing.

The model’s primary limitations are its biophysical and computational assumptions. We modeled cortical columns as single dipoles—a computationally efficient simplification that is reasonable for LFPs (Kajikawa & Schroeder, 2011; Nunez et al., 2019). However, a more realistic model would incorporate ROI-specific cell types, laminar structures, cortical thickness, and multiple biophysical signal mixing. Furthermore, our information model ignores the mutual information across the array by computing information from the single highest SNR channel and thus underestimates some information in the array.

Computationally, we prioritized flexibility and efficiency by assuming an isotropic medium for the lead field. This assumption eliminates the need to recompute the lead field for each device position, reducing processing time from weeks to hours. Although tissue anisotropy from structures such as white matter or CSF is not modeled, the approximation remains reasonable for mesoscale gray matter sources, which are largely isotropic. Distant sources more likely to be affected will also be much lower in measured magnitude and may also lie outside of the 5-10 mm lead field’s primary range. For applications targeting a specific structure, a more tenable method exists. If device placement is defined first, the lead field can be computed for a fixed set of sources, reducing computation while permitting a more accurate anisotropic model. However, this strategy is practical only for small ROIs with few sources, where placement is likely to be more naïve.

### 4.2. Multi-device trajectory optimization significantly improves information capacity

Surgery planning for intracranial implants, SEEG or DBS, is currently performed by a surgeon’s trajectory decisions avoiding blood vessels, vesicles, sulcal entry points, and white matter location. This method reduces complication rates but does not aim to improve recording quality. The method we present is meant to serve as a secondary, complementary step to enhance recording quality.

Genetic optimization applied in §3.3 shows consistent results by measure of total information capacity in the source domain. The information capacity of the system increases with diminishing returns as expected due to the measurement of mutual information within the ROI. It should be noted that this test did not consider signal montaging which would improve SNR of some sources. Solution initialization exceeded human selection, but these methods may prove more valuable as a tool for trajectory refinement given clinician or researcher input.

This method’s agnostic nature toward devices and sources used in optimization offers flexibility by using a variety of device types and their unique lead fields. Additionally, individual device limitations such as depth, angle, and proximity allow for realistic arrangements to be generated. The source space provides similar flexibility in that an anatomical MRI scan can be used. The value of trajectory optimization is most likely to be seen when inter-device montaging is used.

In addition to the limitations mentioned regarding simulation, there are limitations specific to the MRI source model and optimization problem. BrainSuite (Shattuck & Leahy, 2002) is used exclusively for cortical surface extractions and automatic segmentation. Though stable, there is a level of assumed error with image to volume conversion and in segmentation. In computation, the inner cortical surface serves as our primary reference with a defined offset to the fourth layer of cortex. A key point for future improvement is to use local curvature of cortex in the calculation for layer four offset and likewise adjusting the dipole strength. Though the optimization solutions are successful relative to human efforts, it is extremely unlikely that they ever find the global maximum due to the non-convex nature of this problem, particularly with multiple devices. This is an NP-hard problem and an example of the P versus NP problem, where individual solutions are quick to test but it is practically impossible to find the best global solution or test all possible solutions.

Prospective development will benefit from more accurate reconstruction of T1-MRI cortical surfaces and curvature corrected sources. Additionally, delicate structure avoidance would provide the benefit of existing surgical planning in addition to these enhancements. Our current research is underway to increase the dipole model detail and add specific characteristics unique to different cortical regions. Evaluation of different optimization methods is also in progress to further enhance results and reduce computational load.

### 4.3. SEPIO enables classification with sparse sensor subsets

Sparse sampling, or compressed sensing, is a common technique in signal analysis in many fields and taking many forms, originating from the 𝐿^1^ − 𝑛𝑜𝑟𝑚 of Laplace. More recently, Brunton *et al*. explored sparse sensing applied to standard and novel classification tasks (Brunton et al., 2016). Notably, this included selection of limited sensors based on complete datasets as well as selection based on subsampled datasets, both of which are capable of reproducing successful classification given sufficient sampling of the signals’ inherent low-dimensional patterns found in principal component space. In this study, sparse sensing is applied to sensor voltage in order to investigate the potential for reduced sensing in electrophysiology and further refinement of the trajectory. Though the optimization of trajectory discussed prior is a non-convex problem, SEPIO may allow a given trajectory to be considered convex for the usage of these methods in sensor sorting. (Brunton et al., 2016)

In order to properly evaluate this method, *in silico* data are required for supervised training with well-defined classes. (Næss et al., 2021). The two-sulcus model (§3.5) utilizes 128 source dipoles with up to two devices, an intracortical and a subdural grid. Classification with SEPIO is moderately successful with any individual device but is improved greatly when considering both an intracortical probe and subdural grid. In this case, the sensor order can be mixed between two different source perspectives and gains accuracy faster for each sensor addition. The best case uses a DiSc and ECoG, where the reduced set to 146 sensors achieves over 90% accuracy for the 128-class system. This level of success with great spatial variety suggests that fewer sensors may be necessary with an advanced recording plan that achieves greater diversity in lead field montaging.

The utility of simulation in early testing is undeniable but requires real-world comparison. The single sulcus phantom model (§3.6) demonstrates a similar example using two intracortical DiSc devices and 11 dipoles combined in choose-2 fashion to form 55 trial classes. Both simulation and phantom datasets achieve accuracy over 95% with all 128 sensors for the 55-class system. More interestingly, the top 16 sensors selected from DiSc (128 total) achieve 75% and 83% accuracy for simulation and phantom, respectively. A basic 4×2 array at the top and bottom of both devices reached 59% and 55%, respectively. Virtual macros fall much farther behind due to signal averaging over the large contacts. Swapped SEPIO selections for simulation and phantom maintained a high level of accuracy suggesting that the two datasets are similar enough that solutions are interchangeable. Interestingly, some points of the swapped sensor selections exceed the full dataset accuracy but drop below at the maximum sensor count. Swapping results seem to serve as a method to explore other high-quality results for intermediate sensor counts in addition to validating the quality of *in silico* reconstruction of the phantom.

Simulation and phantom assessments provide the initial evidence to suggest that SEPIO is effective in sorting sensors in priority order for electrophysiology classification. Limitations include a high sensor to source ratio (SSR) that leads to easier classification by SEPIO and may cause ‘lazy’ sorting of sensor priority. On the other hand, when sparse sensing occurs then the classification is more difficult, and the optimization effort is most impactful. Broadly speaking, one can assume that epilepsy diagnosis would have few seizure sources of large SNR relative to other background signal. Our phantom model also had few sources relative to sensor count. In the phantom model, the moderate differences in SEPIO classification accuracy between different electrode patterns suggest that 16 sensors is within a reasonable range to improve results. BCI applications would have many more potential sources and information states, making a more valuable opportunity to improve sensing in a low SSR environment.

Sparse sensor classification is a useful tool, but surgical inaccuracy may account for greater spatial variation than the width of intracortical devices (Vakharia et al., 2017), reducing effective benefit from precise placement. Additionally, subject-specific MRI data is likely available to construct a tissue model for dipole locations but key characteristics of charge and length, must still be assumed (Murakami & Okada, 2015).

## 5. Conclusion

We demonstrate that genetic optimization of device trajectories achieves significant increases in the total information capacity objective function versus human selections, and results in high ROI coverage. SEPIO applied to optimized trajectories provides confirmation that classification accuracy increases with device and sensor count and provides a sorting order for further refinement. This combined method gives a quantitative measure to directly compare a variety of recording devices, finding synergistic combinations unique to each ROI. The goal is that this tool will be used by others to provide a metric for technology selection and application. Optimization methods identify key implantation sites while SEPIO enhances sparse sensor sets, with the potential to reduce the number of probes required. We anticipate the strongest impact will be in situations where ROI coverage is critical, source reconstruction is desired, sensor counts are limited, customization is possible such as with additive manufacturing technologies, and especially in humans when a pre-surgery structural MRI is provided. Improvements in implantation may then aid surgeons and reduce surgical burden on patients.

This versatile approach to refining trajectory plans by predicting information capacity is an important alternative to iterative improvements in animal experiments. Improvements in trajectory optimization requires further refinement of source space models as well as an expanded library of device lead fields, allowing ease of comparative multi-device studies. Altogether, this first step presents a miscellany of directions worthy of investigation and demonstrates the value of trajectory optimization and SEPIO for improvement and quantitative comparisons of devices and arrangements in a subject-specific simulation.

## Acknowledgements and Contributions

JAW, CEW, AMA, NT, and JPS received funding from NIH NINDS UG3NS125487, NIH NINDS RF1NS133972, and Rice IITK global initiative funding.

CES received funding from NIH NINDS RF1NS133972, P50 MH109429, and R01 DC102947

JCM received funding from NIH/NIBIB R01EB026299, NIH NINDS UG3NS125487, and NIH NINDS RF1NS133972.

AAJ and RML received funding from NIH/NINDS R01NS121761, NIH/NIBIB R01EB026299, and DOD/ USAMRAA HT94252310149.

YR received support from the Rice SURF program.

The following are author contributions to this work using the CRediT (Contributor Roles Taxonomy) system. All authors contributed to draft review and editing. JAW contributed conceptualization, methodology, software, formal analysis, supervision, and original draft writing. CEW contributed conceptualization, methodology, software, formal analysis, and original draft writing. RZ, YR, and JS contributed conceptualization, methodology, and software. AMA contributed conceptualization and methodology. TM contributed conceptualization and methodology. PM contributed methodology. JCM contributed conceptualization, methodology, and software. AR contributed methodology and resources. CES contributed resources. AAJ and RML contributed software and resources. NT contributed resources and funding acquisition. JPS contributed conceptualization, methodology, formal analysis, supervision, original draft writing, resources, funding acquisition, and project administration.

## 6. Ethical statement & Data Availability

No ethical review was necessary as all MRI data was acquired from Beijing Normal University (CC-BY-NC) under the International Neuroimaging Datasharing Initiative (INDI), *via* BrainSuite.

There are no conflicts of interest to disclose.

Code is publicly available on the GitHub repository TBBL-UTHealth/SEPIO (Translational Biomimetic Bioelectronics Laboratory, 2025) with required data referenced in Zenodo repository.

## Supplemental Data

**Supplemental Figure 1.**
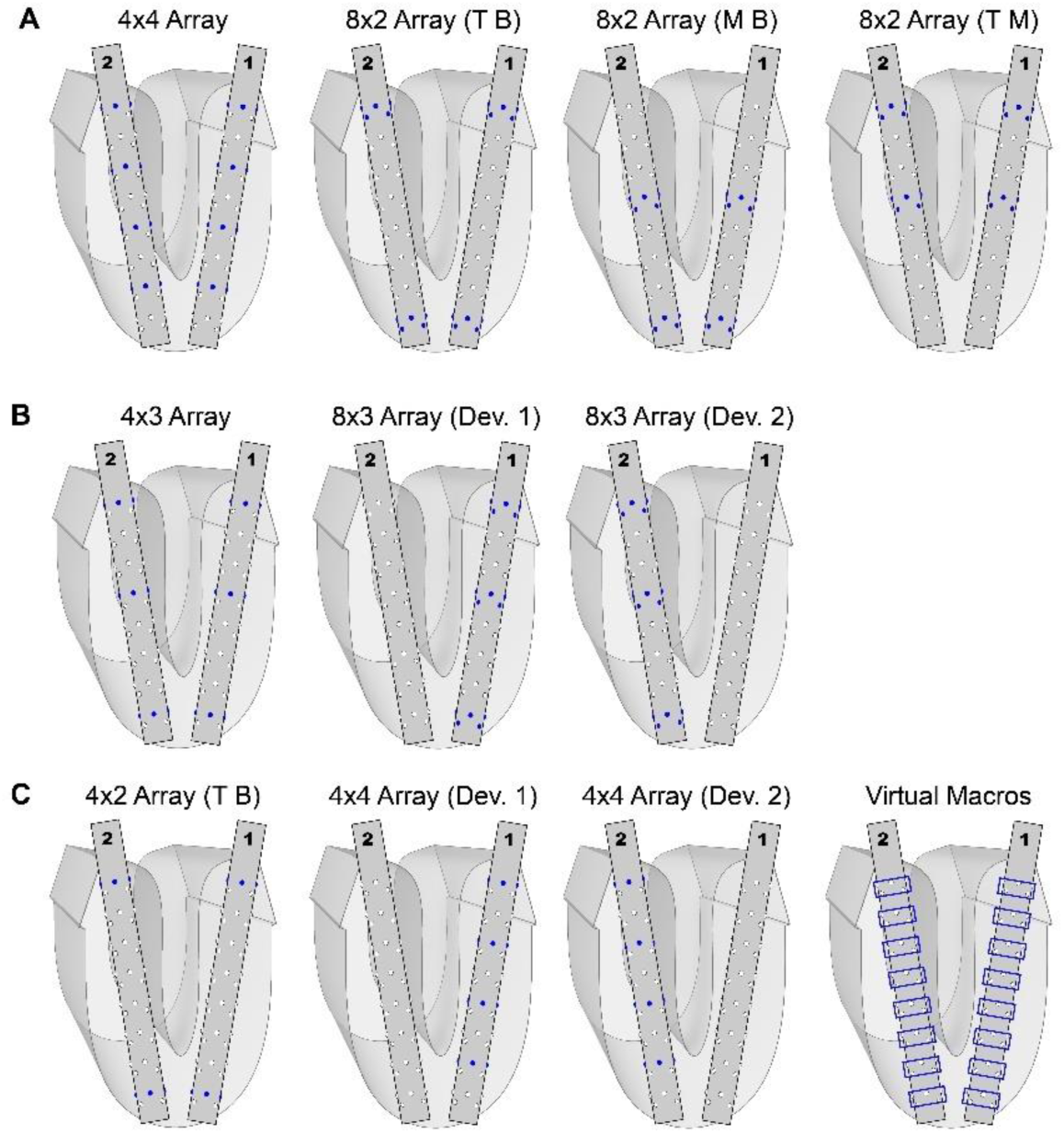
Demonstration of standard array arrangements using two DiSc devices. By row, these devices contain (A) 32, (B) 24, or (C) 16 active sensors. Classification results from these arrangements are given in Supplemental Table 1.

**Supplemental Figure 2.**
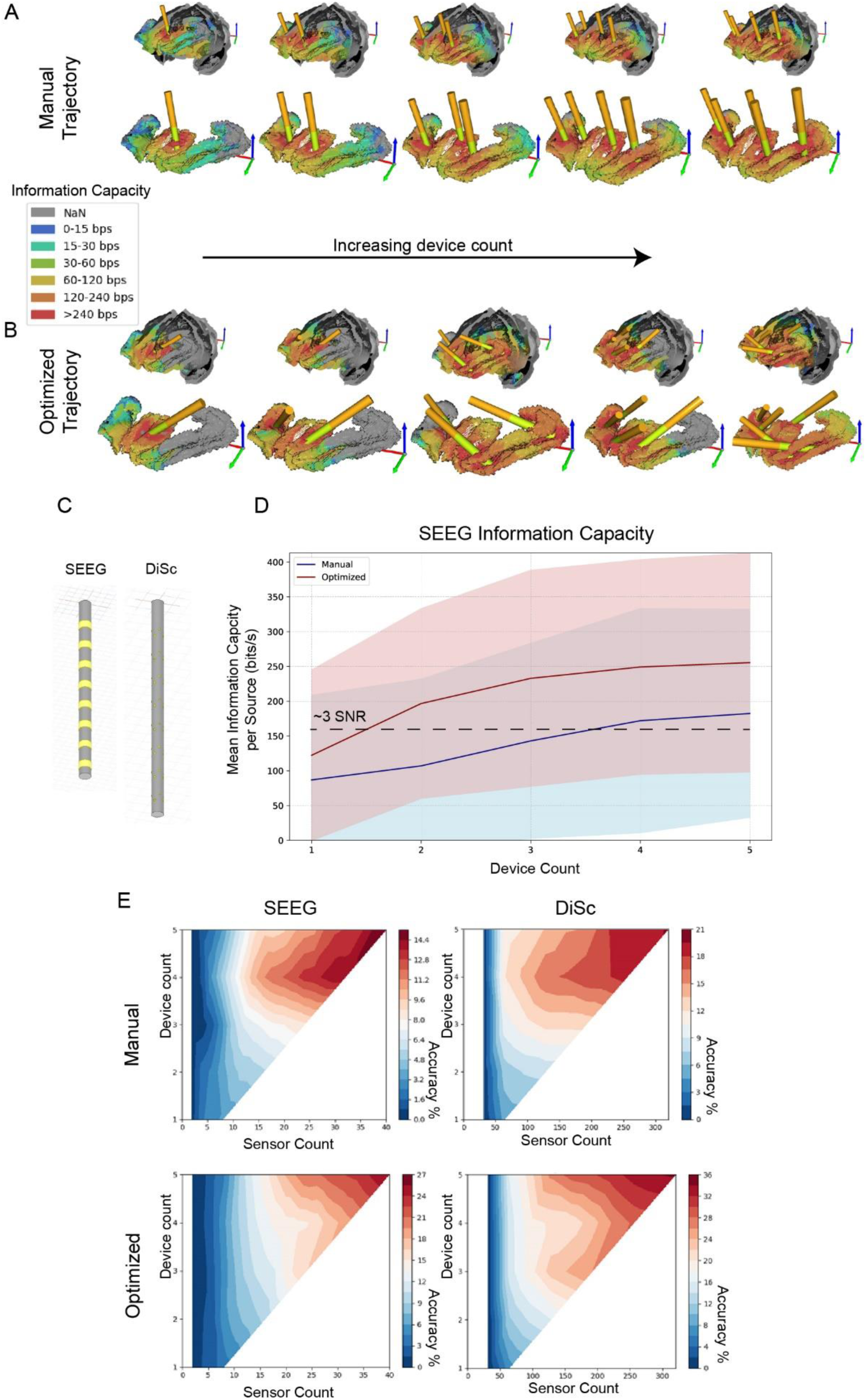
Case study in macaque anterior cingulate cortex (ACC) and dorsolateral prefrontal cortex (dlPFC) using SEEG and DiSc. (A&B) Bilateral ACC+dlPFC trajectories are visualized with an information capacity heatmap. One to five 8-channel SEEG are shown with the recording probe (yellow) and rigid backend (orange) visualized relative to the ROI and whole cortex. (A) Manual trajectories were placed similar to published literature while (B) optimized trajectories were derived from our software. A heatmap of information capacity depicts the maximum potential recording value for each point on the cortex. Training was performed with the ACC weighed twice (per source) that of the dlPFC to encourage greater central coverage. (C) The two device models have a comparable profile and total span, viewed in ANSYS. (D) Mean information capacity across the ACC+dlPFC ROI for each device count in mean and optimized trajectories. Values are calculated on the optimization test dataset. Mean lines are shown with shaded regions depicting one standard deviation. A dashed line is added at a level depicting roughly 3 SNR signal acquisition. (E) SEPIO heatmaps comparing trajectories and devices. Heat map range and sensor count varies between SEEG and DiSc. Percent accuracy is determined from SEPIO classification accuracy on test data withheld during model training. 13,658 unique sources and classes yield chance accuracy below 0.01%. **Notes on Supp.** Fig. 2: This ACC-dlPFC ROI provides a much larger, diverse, and difficult testing environment for trajectory optimization. Device orientation diversity improves the potential signal measurement and diversity when source orientations vary. This is observed in all optimized trajectories with two or more devices. Often, these inter-device angles vary by 20° or more, occasionally becoming nearly orthogonal. Since this algorithm considers device spacing and potential collision of the device backend, the distribution tends to follow directional trends with clusters of devices pointing in similar directions to improve packing density, particularly for smaller ROI such as Broca’s area seen previously. With five devices in the ACC-dlPFC, manual trajectory reached a mean of 182 bit/s while optimized reached 256 bits/s, a 40.7% improvement. Classification by SEPIO provides further insight into downstream potential for the trajectories. In manual trajectories, SEEG achieved a maximum accuracy of 14.9% and DiSc achieved 19.6%. Accuracy is understandably low given that the dlPFC contains 10,290 sources, totaling over four times ACC alone. In optimized trajectories, SEEG achieved a maximum accuracy of 25.6% and DiSc achieved 34.3%. Through optimization, classification improved by 4.7 points for SEEG and 15 points for DiSc.

**Supplemental Table 1.** SEPIO accuracy results for several default array designs visualized in supplemental figure 1. In these comparisons, 32, 24, or 16 sensors are allowed for either standard arrays or as a limit for SEPIO to choose from the total 128 sensors across two DiSc. Percentile accuracy is given with parentheticals containing the change in percentile accuracy relative to the accuracy of SEPIO. Simulation and phantom are kept as separate datasets. All array accuracies are done in Monte-Carlo fashion. Bold and underlined values are displayed in Figure 14.

| **32 Sensors** | SEPIO | 4x4 Array | 8x2 Array,<br>T B | 8x2 Array,<br>M B | 8x2 Array,<br>T M |
| --- | --- | --- | --- | --- | --- |
| Simulation Accuracy: | <u>84%</u> | <u>83% (-1%)</u> | 71% (-13%) | 70% (-14%) | 80% (-4%) |
| Phantom Accuracy: | <u>97%</u> | <u>93% (-4%)</u> | 70% (-27%) | 82% (-15%) | 95% (-2%) |
| **24 Sensors** | SEPIO | 4x3 Array,<br>Both | 8x3 Array,<br>Dev 1 | 8x3 Array,<br>Dev 2 | - |
| Simulation Accuracy: | <u>83%</u> | <u>76% (-7%)</u> | 45% (-38%) | 44% (-39%) | - |
| Phantom Accuracy: | <u>95%</u> | <u>89% (-6%)</u> | 71% (-24%) | 68% (-27%) | - |
| **16 Sensors** | SEPIO | 4x2 Array,<br>T B | 4x4 Array,<br>Dev 1 | 4x4 Array,<br>Dev 2 | **Virtual Macros,**<br>**Both** |
| Simulation Accuracy: | <u>75%</u> | <u>59% (-16%)</u> | 62% (-13%) | 58% (-27%) | <u>14% (-61%)</u> |
| Phantom Accuracy: | <u>83%</u> | <u>55% (-28%)</u> | 71% (-12%) | 73% (-10%) | <u>11% (-72%)</u> |

